# Extensive sex differences at the initiation of genetic recombination

**DOI:** 10.1101/235002

**Authors:** Kevin Brick, Sarah Thibault-Sennett, Fatima Smagulova, Kwan-Wood G. Lam, Yongmei Pu, Florencia Pratto, R. Daniel Camerini-Otero, Galina V. Petukhova

## Abstract

Homologous recombination in meiosis is initiated by programmed DNA double strand breaks (DSBs) and DSB repair as a crossover is essential to prevent chromosomal abnormalities in gametes. Sex differences in recombination have been previously observed by analyses of recombination end-products. To understand when and how sex differences are established, we built genome-wide maps of meiotic DSBs in both male and female mice. We found that most recombination initiates at sex-biased DSB hotspots. Local context, the choice of DSB targeting pathway and sex-specific patterns of DNA methylation give rise to these differences. Sex differences are not limited to the initiation stage, as the rate at which DSBs are repaired as crossovers appears to differ between the sexes in distal regions. This uneven repair patterning may be linked to the higher aneuploidy rate in females. Together, these data demonstrate that sex differences occur early in meiotic recombination.

## Main text

Genetic recombination brings homologous chromosomes together during meiotic prophase and facilitates their orderly segregation at the first meiotic division. Recombination is initiated by programmed DNA double strand breaks (DSBs) that are subsequently repaired as either crossovers (COs) or as non-crossovers (NCOs). The pattern and frequency of recombination can differ between males and females of the same species: the female CO rate is higher in humans and mice^1^ while in most studied mammals COs are highly concentrated at sub-telomeric regions in males, but not in females^2–6^. This pattern is not universal however and in some species, such as pigs, subtelomeric crossovers are elevated in females^7^. To date, sex differences in recombination have been studied by comparing the genetic end products of recombination, primarily COs, between the sexes. However, CO-based estimates of recombination rates are limited by the number of sampled meioses and the maximum resolution of such analyses is determined by the spacing of sequence polymorphisms. Most importantly, sex-specific variation that manifests at the initiation of meiotic recombination has not been studied and it remains an open question whether sex differences originate at the time of DSB formation or whether they arise later, during DSB repair. We therefore generated quantitative, high resolution and genome-wide maps of meiotic DSBs in both male and female mice to examine when and where sex biases in recombination are established, and to elucidate the mechanism(s) that give rise to these biases.

To map meiotic DSBs in female meiosis we exploited a method we previously developed^8^ to map DSB hotspots in mouse^9–11^ and human^12^ males. This variant of ChIP-Seq (single stranded DNA sequencing, SSDS) detects single stranded DNA (ssDNA) bound to the DMC1 protein, an early intermediate in the DSB repair process^12,13^. In female mice, meiotic DSBs form in the fetal ovary and the number of cells undergoing DSB repair are maximal between 15 and 16 days post coitum ^14^ (63-84% of meiocytes are in leptotene/zygotene stages; approximately 10,000 cells per ovary^14^). Each fetal ovary has 100 times fewer such cells than in adult testis, therefore mapping DSBs in individual ovaries is not feasible. Instead, we harvested ovaries at 15.5 dpc to generate one pool of 90 and one pool of 230 fetal ovaries for DSB mapping. From the 230-ovary pool, we generated a DSB map of similar quality to that of nine independent DSB maps generated from male individuals (Fraction of Reads in Peaks (FRiP) = 33%; FRiP for testis maps = 22-47%; Figure 1A; Sample O1, Supplementary Data Figure 1, 2A). The DSB map from the 90-ovary pool was of lower quality (FRiP = 6%; Supplementary Data Figure 1B) but shared most hotspots (6,765 / 7,233; 94%) with the better ovary DSB map (Sample O2, Supplementary Data Figure 1). These data provide the first maps of meiotic DSBs in mammalian oocytes.

**Figure 1.**
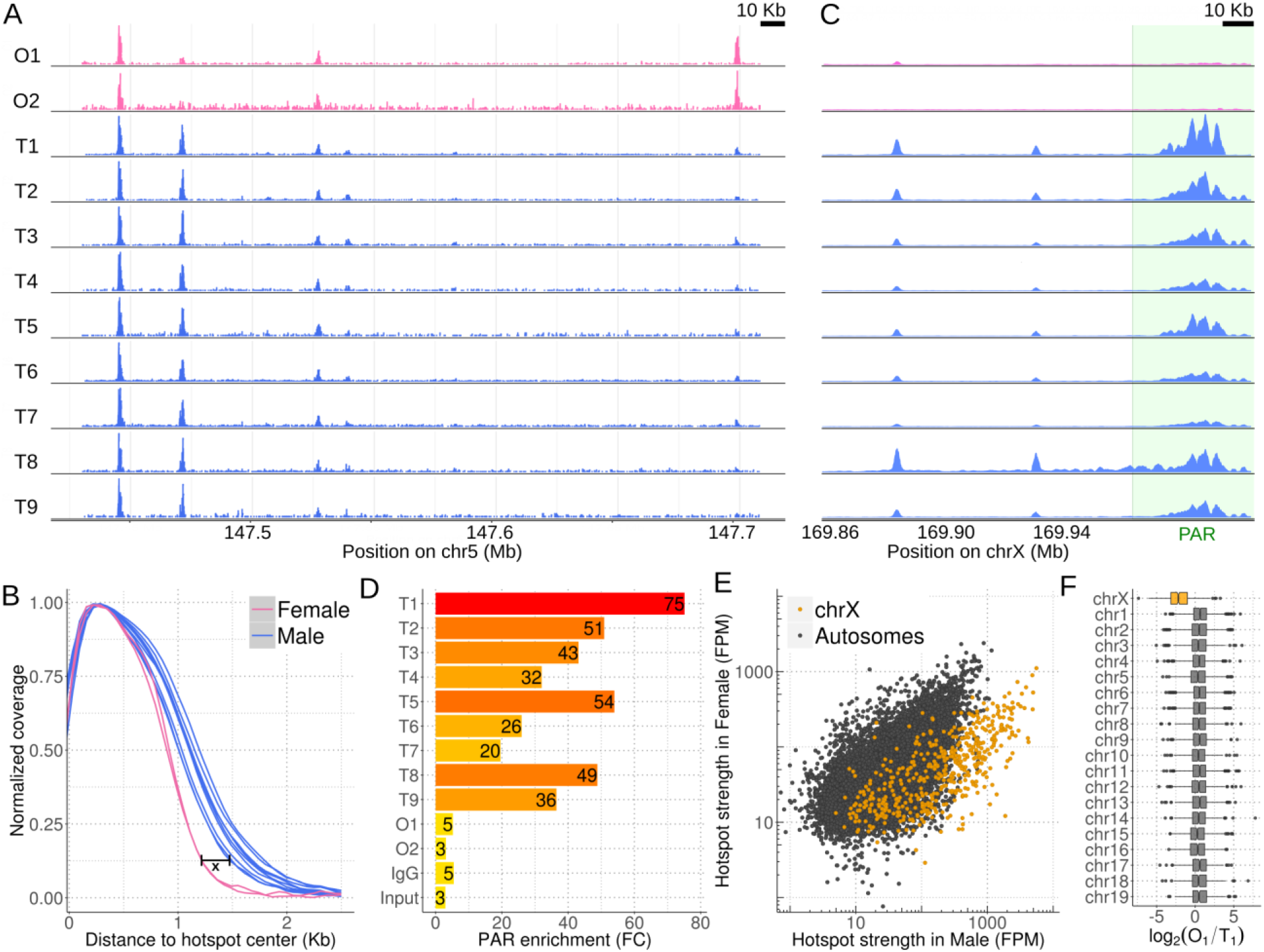
Meiotic DSB hotspots can be detected in female mice. (A) DSB maps from ovaries (O1, O2) and from testis (T1-9). (B) SSDS coverage at DSB hotspots is wider in male than in female mice. Coverage is plotted as the 3’ distance along ssDNA to the hotspot center. Male samples are shown as blue lines, female as pink. The interval x is the widest point between the distributions (~400bp). (C, D) PAR SSDS signal is enriched in male, but not in female meiosis. (C) Coverage is normalized by ssDNA fragments on chrX. The ~50 kb of the mouse PAR adjacent to the pseudo-autosomal boundary (green box) is the only assembled PAR sequence in the genome. (D) Enrichment is the 50 Kb PAR SSDS signal divided by the mean coverage for all chrX 50 Kb intervals; fold change (FC). All male libraries show enrichment relative to controls, whereas female samples do not. The PAR is also enriched relative to other regions in input and IgG samples, likely because the mouse PAR is highly repetitive and poorly assembled. (E, F) DSB hotspots on chromosome X appear stronger in male meiosis. (E) Strength is shown as SSDS fragments per million (FPM). (F) Hotspots in females appear weak relative to males on chromosome X compared to all autosomes.

Most DSB hotspots are found in both sexes (Supplementary Data Figure 2A); 88% of hotspots from the better ovary DSB map are found in males and this increases to 97% of hotspots common to both ovary maps. Hotspots unique to either sex are weak (Supplementary Data Figure 2B,C) and contribute < 2% of the SSDS signal. Given that strength estimates at weak hotspots are noisy and that ChIP-Seq provides relative rather than absolute estimates of hotspot use, it is likely that these hotspots are also used in males, but with a frequency below our detection threshold. Hotspots found only in males are also relatively weak (Supplementary Data Figure 2B,D). Thus, we conclude that there are few, if any, hotspots that are used exclusively in either sex. One intriguing difference between the sexes, is that the DMC1-SSDS signal at hotspots is narrower in females than in males (by ~400 bp at the widest point; Figure 1B, Supplementary Data Figure 3A-D). This may result from shorter resection around DSBs in females, from DMC1 loading over a shorter distance in females or from different repair dynamics between the sexes (Supplementary Data Figure 3E). Irrespective of the mechanism, these data add to a growing body of evidence^15–18^ that DSB processing differs substantially between the male and female germ line.

There are striking differences in meiotic DSB repair on the sex chromosomes in males and females. Since the X and Y chromosomes only share approximately 700 kb of homology (in the pseudoautosomal region, PAR)^19^, an obligate meiotic crossover at the PAR is required in males, the heterogametic sex. Females have two copies of chrX, therefore crossover formation in the PAR is not essential. We found that DSBs in the PAR were highly enriched in all nine males relative to controls, but in neither female (Figure 1C,D). This is consistent with the PAR CO rate being 7-fold higher in males compared to females^20^. The dynamics of DSB repair on chrX outside the PAR also differs between the sexes. On the single male copy of chrX, DSBs either remain unrepaired^21^ or are continually formed^22^ long after autosomal DSBs have been repaired. This prolonged presence of DSBs on chrX biases their detection, such that in wild-type male mice, hotspots on chrX appear disproportionately strong relative to autosomal hotspots. Indeed, we found that compared to autosomal hotspots, hotspots on chrX appear stronger in males than in females (Figure 1E,F). Because of these systematic differences, the sex chromosomes are excluded from subsequent analyses unless explicitly mentioned. Together, we conclude that sex differences are apparent in DSB maps of chromosome X and that the PAR is not particularly recombinogenic in female meiosis.

The SSDS signal at hotspots is highly reproducible in males, with little inter-individual variability in hotspot usage among the 9 males studied (Figure 2A,C; Spearman R^2^ ≥ 0.90). Similarly, the SSDS signal at hotspots is highly correlated between the two female DSB maps, albeit to a slightly lesser degree than for males (Figure 2B; Spearman R^2^ = 0.78). Unlike the male maps, the female maps were generated from pools of multiple individuals, therefore this small difference may reflect variance in our estimate of the population mean resulting from stochasticity of hotspot targeting in individual females. This seems unlikely however, as the mice are genetically homogeneous and we see negligible inter-individual variation in males. Alternatively, noise in SSDS estimates for the lower quality map simply reduces the correlation. Importantly, the SSDS signal at hotspots is strikingly different between males and females (Figure 2C,D; Spearman R^2^ ≤ 0.33). Indeed, examination of all DSB hotspots found in the better male (T1) or better female (O1) sample (20,119 hotspots; see methods) revealed that 48% of autosomal hotspots are sex-biased (P < 0.001, MAnorm^23^ - see methods; Figure 2D; N_male-bias_ = 4,169 [22%], N_female-bias_ = 5,021 [26%]; N_unbiased_ = 9,863 [52%]). The average sex-biased hotspot differed between the sexes by 4.0 ± 4.3 fold (mean ± s.d.; median = 2.7 fold), and 1,746 hotspots showed over 5-fold difference (Supplementary Data Figure 4A). Sex-biased hotspots are likely under-detected, because at stronger hotspots, where we have the greatest power to detect sex differences, over 60% of hotspots are sex-biased (Supplementary Data Figure 4B). Importantly, sex biases are consistent between the O1 and O2 samples (Fig. 2E, Supplementary Data Figure 5), therefore they reflect true sex differences, and not sampling noise in the pooled ovary maps (see above). 44 ± 0.4 % of the SSDS signal in males and 51 ± 4 % (mean ± s.d.) in females occurs at hotspots biased to their respective sex (Supplementary Data Figure 4C). In addition, a further 16 – 21% occurs at hotspots biased to the other sex (female-biased hotspots in males or male-biased hotspots in females). Therefore, although most hotspots are used in both sexes, the majority of the SSDS signal, in both sexes, originates at sex biased hotspots.

**Figure 2.**
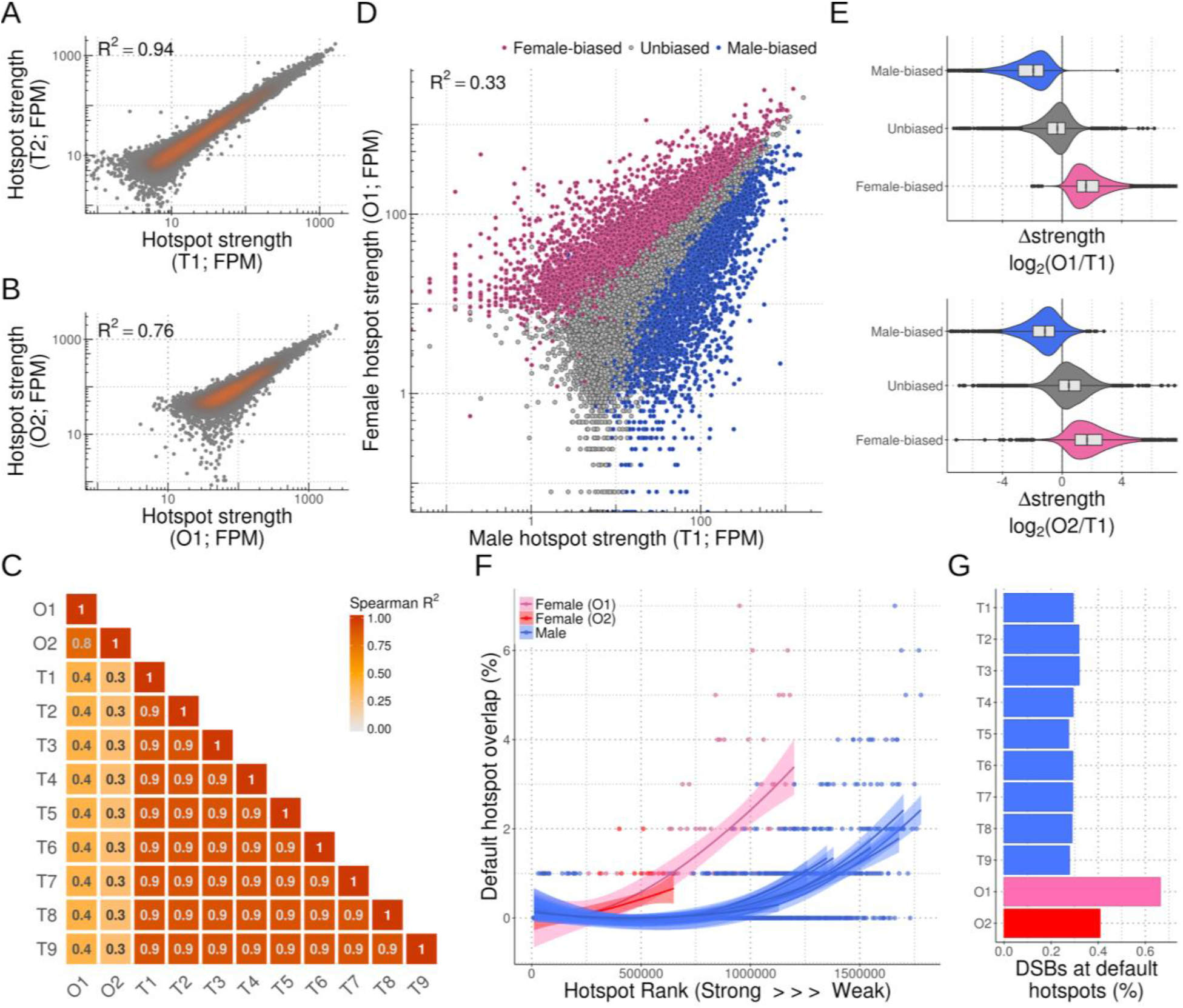
Extensive sex differences in DSB hotspot usage. Hotspot strength is highly correlated between (A) male DSB maps and (B) female DSB maps. Strength for each sample is expressed in fragments per million (FPM). Only hotspots called in both maps are considered. (C) Correlation matrix illustrating consistent SSDS strength estimates between maps derived from the same sex, but differences between males and females. (D) The SSDS signal at hotspots differs substantially between males and females. Female-biased (pink), unbiased (grey) and male-biased hotspots (blue) are shown. (E) Sex-biased hotspots show similar biases in the O1 and O2 DSB maps. (F) Default hotspots are used more frequently in female meiosis. Hotspots identified in each library were ranked by strength and split into 250 hotspot bins. Hotspots that overlap hotspots in a Prdm9^−/−^ mouse but that lack a putative PrBS were designated as “default” hotspots for each strength-ranked bin. In the female samples, default hotspots were relatively strong. (G) More DSBs occur at default hotspots in female meiosis than in male meiosis.

SSDS is an accurate measure of DSB frequency (hotspot strength), tightly correlated with an independent measure of hotspot strength in male mice^21^. Nonetheless, since the SSDS signal is likely affected by the lifespan of DSB repair intermediates^21,24^, a component of the observed sex biases may arise from differential DSB repair dynamics between the sexes. To establish if sex biases precede DSB formation and repair, we examined Histone 3 Lysine 4 trimethylation (H3K4me3), a histone modification introduced at hotspots by PRDM9 that is thought to recruit the DSB machinery^25^. The H3K4me3 signal at hotspots correlated better with SSDS signal from the respective sex (Supplementary Data Figure 6): 69% of female-biased hotspots coincided with a H3K4me3 ChIP-Seq peak in fetal ovary, but just 39% of male-biased, and 43% of unbiased hotspots overlapped these sites. Importantly, the sex-biased hotspots defined using SSDS showed similar sex biases in H3K4me3 signal (Supplementary Data Figure 6). The magnitude of sex bias is reduced in H3K4me3 ChIP-Seq compared to SSDS, perhaps a reflection of the reduced sensitivity of H3K4me3 ChIP-Seq at hotspots compared to SSDS. Thus, while the contribution of repair dynamics to sex biases remains unclear, sex biases in recombination are established before DSB formation.

PRDM9 targets the vast majority of DSBs to hotspots^9^ through DNA sequence-specific binding. In the absence of PRDM9, a default and perhaps ancient mechanism targets DSBs to hotspots at PRDM9-independent H3K4me3 modified nucleosomes^9^. Although PRDM9 defines the majority of DSB hotspots in both sexes we found that the default targeting pathway is used far more frequently by females (pink & red lines; Figure 2F) than by males (blue lines). Nevertheless, default hotspots are weak, contributing 0.4 – 0.6% of the SSDS signal in females but just 0.2% in males (Figure 2G). Since default hotspots account for 8.5% (427/5,021) of female-biased hotspots but only 1.3% (58/4,169) of male-biased hotspots, we distinguish between hotspots defined by each pathway for subsequent analyses. Default hotspots are used despite an abundance of PRDM9 binding sites (PrBSs)^26^, which may imply that either PRDM9 or some other component of the DSB machinery is limiting in oocytes. Alternatively, since default hotspots coincide with functional genomic elements, including transcription start sites (TSSs)^9^, the DNA at these sites may be more accessible in female meiosis. Intriguingly, recombination (CO rate) also increases locally around TSSs in human females, but not in males^27^. Although this has been proposed to be a PRDM9-dependent increase, it remains to be seen if the default targeting pathway is also active in humans^28^.

In addition to primary DNA sequence, the chromatin context can modulate the usage of DSB hotspots; for example, PRDM9-independent H3K4me3 modified nucleosomes and heterochromatic histone modifications disfavor hotspot usage^24,26^, whereas the SSDS signal is elevated in actively transcribed regions^11,26^. Since meiotic chromosome packaging differs between males and females^29^ we looked for evidence of large-scale epigenetic effects that modulate sex biases. Both male-biased and female-biased hotspots occurred in clusters more frequently than expected from random simulations (see methods; Figure 3A-C). These biased hotspot domains are relatively evenly distributed across chromosomes (Supplementary Data Figure 6), and unbiased hotspots did not cluster (Figure 3B,C). Cluster size scaled with the number of hotspots per cluster and we found no evidence for a physical limit on cluster size (Supplementary Data Figure 6). A similar proportion of default and PRDM9-defined hotspots were found in clusters (Supplementary Data Figure 6) suggesting that domain-scale regulation of hotspot usage is independent of the mechanism that targets DSBs. Overall, sex biases in hotspot usage appear to be partly governed by aspects of local chromatin structure and this clustering is consistent with the known spatial clustering of sex biased crossovers in humans^30^. Given our findings, these sex differences in humans may be established at the initiation of recombination.

**Figure 3.**
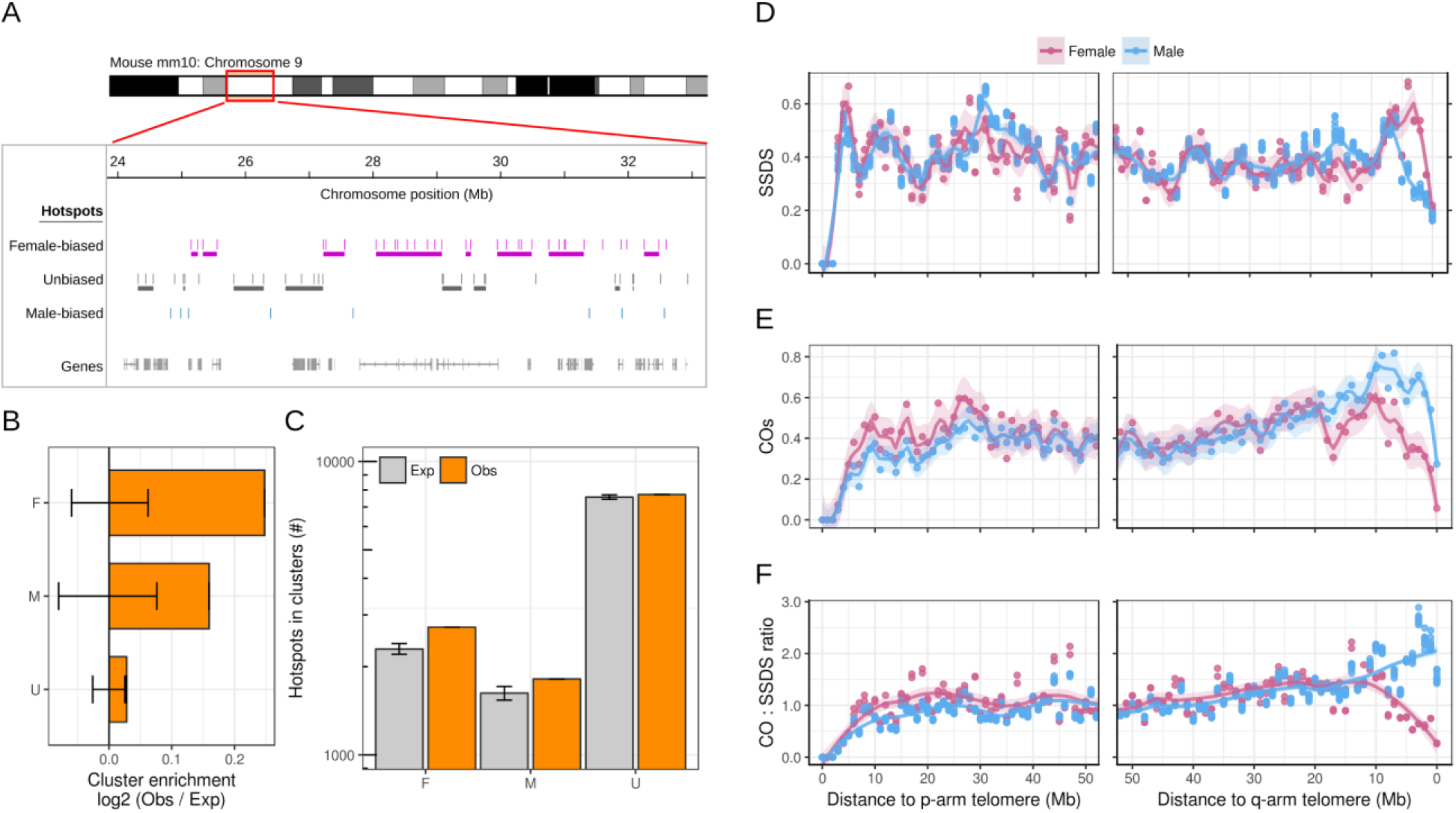
Large-scale influences on sex-biased recombination. (A) Several clusters of female-biased hotspots. Clusters are indicated as horizontal lines below the hotspot tracks. (B) Sex-biased hotspots cluster more than expected in the genome (empirical P < 0.001). A cluster is >1 consecutive hotspots with the same sex bias. The expected cluster counts were calculated by randomly shuffling the sex-biases of hotspots and counting clusters (error bars = ± 99.9% bootstrapped CI). (C) The number of hotspots in clusters is shown. Grey bars represent the median of 10,000 bootstrapped randomized sets (error bars: ± 99.9% CI). (D) Male and female DSB maps deviate adjacent to the q-arm telomere. The SSDS signal decreases in all males (blue) relative to females (pink) in this region. We focus on the sub-telomeric region of the q-arm because unassembled centromeric DNA abuts the p-arm telomere in mice. (E) Genetic crossovers^33^ also exhibit sex biases adjacent to the non-centromeric telomere. Unlike the SSDS signal, crossovers in this region are elevated in males as compared to in females. (F) Sex dimorphism in the CO:SSDS ratio implies that sub-telomeric DSBs are repaired differently in male and female mice. All profiles are generated from 1Mb non-overlapping windows.

Recombination in subtelomeric regions is of particular interest in the context of female meiosis, because in human oocytes, the chromosomes that have a single telomere-adjacent crossover appear to have an elevated risk of mis-segregation^31,32^. This is likely because telomeric chiasmata are less stable through the long dictyate arrest of female oocytes^32^. We found that the SSDS signal in sub-telomeric regions contrasts starkly with that of crossovers: the SSDS signal is high in females relative to males (Figure 3C), whereas, crossovers are less frequent in females (Figure 3D)^33^. Distal crossovers in females may reduce gamete fitness and be under-detected in pedigree studies. However, a recent study that examined repair outcomes in fetal oocytes (where there should be negligible selection against karyotypic defects), also showed crossover depletion at two subtelomeric hotspots in females^34^, consistent with pedigree results. Thus, the crossover : SSDS ratio decreases close to the telomere in females, but increases in males (Figure 3E). In male mice, DSBs appear to be rapidly repaired in the distal 5 Mb, as the SSDS signal is decreased relative to *Spo11*-oligo density (a direct measure of DSB frequency)^35^. It is enticing to speculate that rapid repair of DSBs is a hallmark of crossover designation. This would be consistent with elevated distal CO frequency in males. Alternatively, rapid repair of subtelomeric DSBs may occur in both sexes and differences in SSDS signal may simply reflect differences in the frequency of DSB formation. In either case, sex-specific mechanisms exist to modulate DSB repair outcomes in subtelomeric regions. It will be intriguing to see if these differences are more extreme in humans or other mammals, where dictyate arrest in females can last ten times longer and where selection against distal crossovers may therefore be more pronounced.

A remarkable difference between the sexes at the time of DSB formation is that the genome is globally demethylated in females^36^ but not in males^37^. Since DNA methylation can alter the site preferences of DNA binding proteins^38,39^, we hypothesized that differential DNA methylation may cause sex biases at the initiation of recombination. By examining whole genome bisulfite sequencing data from juvenile testis (13 days post partum (dpp))^40^, we found clear differences in DNA methylation at each set of sex-biased hotspots (Figure 4A; Supplementary Data Figure 7). At male-biased hotspots, the PrBS is frequently methylated, whereas at female-biased hotspots, there is increased DNA methylation in the region adjacent to the PrBS, approximately 75 bp on either side. A methylation “spike” at the PrBS 5’ end is common to all hotspots. Neither sex-specific pattern is seen at unbiased hotspots and both patterns are most pronounced at hotspots exhibiting the greatest magnitude of sex bias (Supplementary Data Figure 8). Meiosis-specific processes do not give rise to these methylation patterns, because qualitatively similar patterns are seen in somatic tissue of both sexes (mammary gland^41^ and male liver^42^), in spermatocyte precursor cells (spermatogonia^43^) and in elutriated pachytene spermatocytes (Supplementary Data Figure 8). Furthermore, these sites do not escape demethylation in the germ line^44^ as these patterns of DNA methylation are absent at early meiotic stages in females (primordial germ cells at 16.5 dpc^36^ (see methods)) (Figure 4A; Supplementary Data Figure 8).

**Figure 4.**
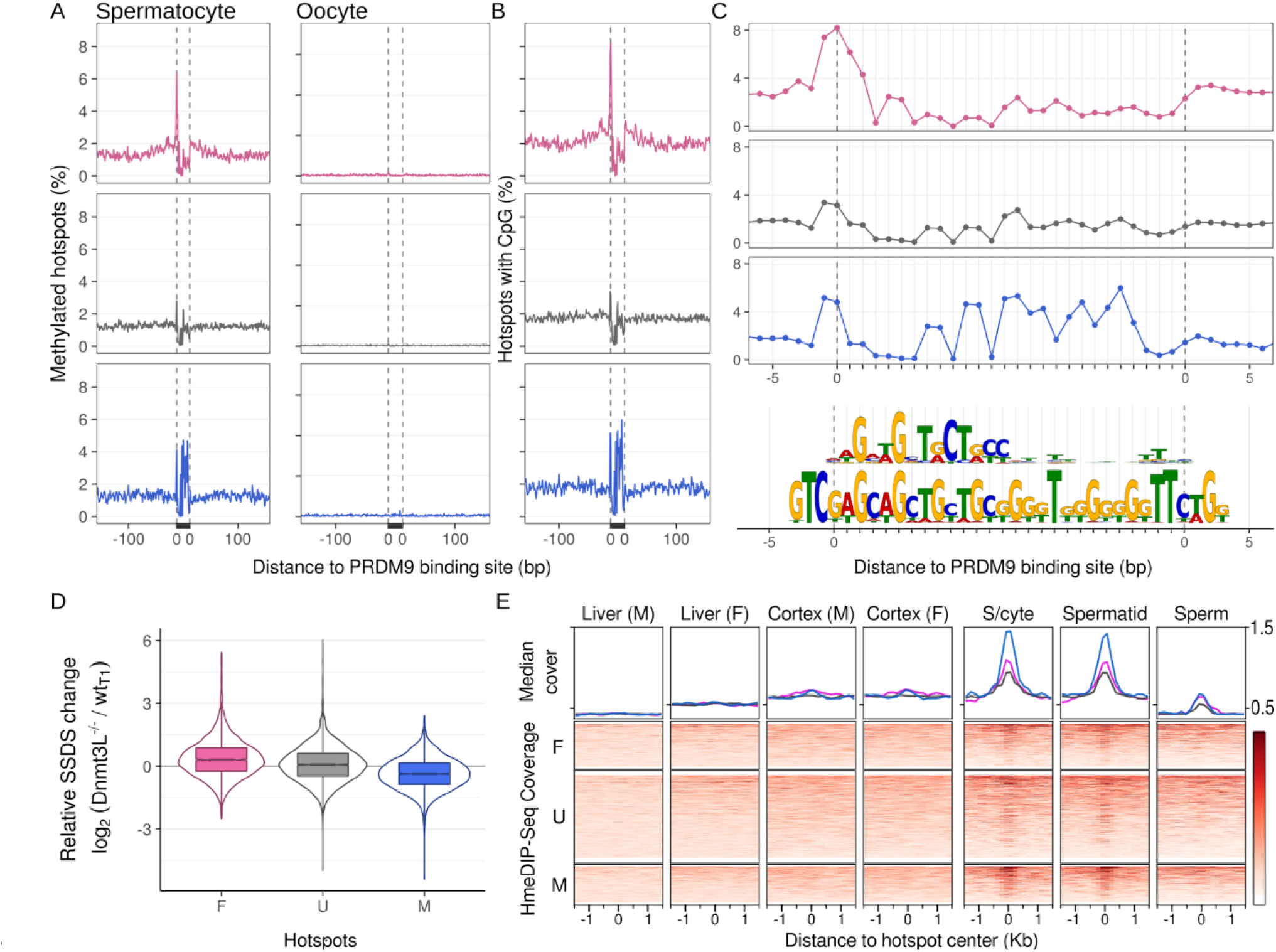
Sex-dimorphic patterns of DNA methylation at sex-biased hotspots. Colors represent female-biased (pink), unbiased (grey) and male-biased (blue) hotspots throughout. (A) Dimorphic DNA methylation patterns at sex-biased hotspots. Left and right panels show average per base methylation from 13 dpp testis^40^ (spermatocyte) or 16.5 dpc primordial germ cells^36^ (oocyte), respectively. (B) DNA methylation patterns mirror the CpG dinucleotide content. (C) CpG density at high resolution around inferred PrBS (upper motif) and the *in silico* predicted PrBS (lower motif). (D) Sex-biased hotspot usage is altered in male mice with reduced DNA methylation (*Dnmt3L^−/−^*). In these males, female-biased hotspot strength is increased (P = 10^−123^, Wilcoxon test) and male-biased hotspot strength is decreased (P = 10^−28^, Wilcoxon test) relative to unbiased hotspots. (E) 5-hydroxymethylcytosine (5-HmC) is enriched at DSB hotspots in the germline, but not in somatic cells. hMeDIP-Seq coverage at hotspot loci in somatic cells (liver & cortex from both sexes^53^), spermatocytes (s/cyte)^43^, spermatids^43^ and sperm^43^ is shown. 5-HmC is particularly enriched at male-biased hotspots in spermatocytes, although female-biased and unbiased hotspots also show enrichment.

The distinct methylation patterns at male- and female-biased hotspots suggest that DNA methylation in males has a dual role in driving sex biases. At PrBS with particular methylated cytosines, PRDM9 binding and DSB formation is favored, whereas, DNA methylation flanking the PrBS disfavors DSBs (Supplementary Data Figure 9). To test this prediction, we generated a DSB map and quantified hotspot strength in male mice that lack *Dnmt3L*, a DNA methyltransferase critical to meiosis^45^. In these mice, DNA methylation is reduced genome-wide^46^. Despite a reported re-localization of DSBs in *Dnmt3L* mice^47^, the vast majority of DSB hotspots identified by SSDS (94 ± 2%; mean ± s.d. in T1-T9) coincide with those in *wt* males. Female-biased hotspots were significantly stronger in male *Dnmt3L^−/−^* mice compared to *wild type* males, whereas male-biased hotspots were weaker (Figure 4B, Supplementary Data Figure 9, 10). This strongly implies that DNA methylation indeed suppresses DSB formation at female-biased hotspots in males, but promotes DSB formation at male-biased hotspots. Importantly, a similar pattern was seen in the *Dnmt3L^−/−^* H3K4me3 ChIP-Seq signal at hotspots (Supplementary Data Figure 10), implying that DNA methylation mediates sex differences before DSB formation. These effects were not quantified as DNA methylation is only partly depleted in *Dnmt3L^−/−^* mice^45^. Together, these data show that DNA methylation plays a dual role in determining hotspot usage: reduced DNA methylation decreases DSB formation at male-biased hotspots but increases DSB formation at female-biased hotspots.

DNA methylation occurs primarily at CpG dinucleotides^48^, and as expected, methylation patterns at hotspots closely reflect CpG density (Fig. 4C). Thus, underlying differences in the DNA bound by PRDM9, by virtue of being frequently methylated, can result in sex biases. At female-biased hotspots, DNA methylation flanking the PrBS appears to suppress hotspot usage. High copy repeats are generally methylated and although they would represent good candidates for a signal of this magnitude we find no repeat elements specifically enriched at female-biased hotspots (see methods). Alternatively, this DNA methylation may favor nucleosome assembly^49,50^ or exert other epigenetic effects that inhibit PRDM9 binding and/or DSB formation at these loci in males. At male-biased hotspots, CpG density is highest at the 3’ end of the PrBS (Fig. 4C; upper motif). This region of the empirically determined PrBS has few apparent binding preferences, however an *in silico* prediction of the PrBS^10,51^ revealed a G-rich consensus at the 3’ end (Fig. 4C; lower motif), consistent with more frequent DNA methylation on the C-rich complementary strand (data not shown). This heretofore unappreciated complexity of PRDM9 binding appears to dictate male sex biases, and therefore, the extent of sex biases may vary for different *Prdm9* alleles. The most prevalent allele of *PRDM9* in humans binds a C-rich sequence and therefore, is likely affected by DNA methylation mediated sex biases. Male-biased recombination in humans is most prominent in distal regions^6,27^ and intriguingly, PrBSs containing a CpG are closer to chromosome ends (28 ± 28 Mb; mean ± s.d.) than other PrBSs (36 ± 29 Mb; mean ± s.d.).

Bisulfite sequencing, the base-pair resolution mapping of methylated DNA, does not distinguish between methylated (5-mC) and hydroxymethylated (5-HmC) cytosines. Therefore, putatively “methylated” nucleotides may be a compound signature of 5-mC and 5-HmC. In contrast to 5-mC (Supplementary Data Figure 8), the 5-HmC signal is absent in somatic cells (Figure 4D). Nonetheless, 5-HmC is highly enriched at DSB hotspots in elutriated (primarily pachytene) spermatocytes^43^ and in spermatids^43^. Enrichment is greatest at male-biased hotspots, with lesser enrichment at unbiased and female-biased hotspots (Figure 4D). Only residual 5-HmC at hotspots remains in sperm^43^, thus, 5-HmC is transiently enriched at hotspots during male meiosis. This implies that 5-HmC or DNA demethylation may contribute to meiotic DSB repair, consistent with previous studies that implicated 5-HmC in the DNA damage response in mitotic cells^52^.

Altogether, these data represent the first comprehensive maps of meiotic DSBs in a female organism and reveal that extensive and heretofore unappreciated sex biases occur at the onset of meiotic recombination. These biases do not linearly translate into differences in the crossover landscape, revealing the multi-layered complexity of sex-biased control of recombination. We have established that several mechanisms modulate sex biases in hotspot formation, including local chromatin structure, choice of targeting and repair pathway, and DNA methylation. Future examination of these mechanisms will yield additional insights into how females and males differentially contribute to shaping evolution of the genome.

## Acknowledgments

We thank P. Hsieh for critical feedback on the manuscript and members of the Camerini-Otero and Petukhova labs for discussion and suggestions. We also thank the NIDDK genomics core for assistance with sequencing. This study used the high performance computational capabilities of the Biowulf Linux cluster at the National Institutes of Health, Bethesda, MD (http://biowulf.nih.gov). This research was supported by NIH grant R01GM084104 from National Institute of General Medical Sciences (G.V.P.), March of Dimes Foundation grant 1-FY13-506 (G.V.P.) and by the NIDDK Intramural Research Program (R.D.C.O.). The sequencing data reported in this paper are archived at the Gene Expression Omnibus (www.ncbi.nlm.nih.gov/geo) as accession no. GSE99921.

## Author Contributions

K.B. performed data analyses and *in-silico* experiments. S.T., F.S., F.P., G.L., Y.P. & K.B. performed wet-lab experiments. K.B. wrote the manuscript. R.D.C-O and G.V.P. supervised the study. All authors contributed to experimental design, discussed the results and critiqued the manuscript.

## Materials and Methods

### Animal procedures

All animal procedures have been approved by the USUHS Institutional Animal Care and Use Committee or were performed according to the NIH Guide for the Care and Use of Laboratory Animals.

### Sample preparation and sequencing

Fetal ovaries were dissected from embryos at 15.5 days post coitum (dpc). Ovaries were dissected in cold PBS and stored at −80°C until use. For SSDS, 90 or 230 ovaries were fixed in 1 ml PBS with 1% paraformaldehyde for 3 min, quenched and homogenized with Dounce homogenizer. Cells were collected by 10 minute centrifugation at 900g using a bucket rotor. The pellet was washed in 1 ml of the following buffers: 1) PBS 2) 0.25% Triton X-100, 10 mM EDTA, 0.5 mM EGTA, 10 mM Tris pH8. Cells were lysed in 0.5 ml of the lysis buffer (1% SDS, 10 mM EDTA, 50 mM TrisCl pH8 with complete protein inhibitor cocktail (Roche)) and the chromatin was sheared with Misonix sonicator with the following parameters: efficiency 1, 10 s on, 20 s off, total sonication time 4 min. Chromatin was cleared by 10 min centrifugation at 12,000g at 4C. The supernatant was diluted 2-fold by ChiP buffer (0.01% SDS, 1.1% Triton X-100, 1.2 mM EDTA, 16.7 mM TrisHCl, 167 mM NaCl) and dialyzed against the same buffer for 5 hour at 4°C.

Chromatin was incubated with 6μg of custom made anti-DMC1 antibody and 20μl Dynabeads (10002D, Invitrogen) at 4°C overnight followed by washing with 500μl of the following buffers: 1) 0.1% SDS, 1% Triton X-100, 2 mM EDTA, 20 mM TrisHCl, 150 mM NaCl; 2) 0.1% SDS, 1% Triton X-100, 2 mM EDTA, 20 mM TrisCl pH8, 500 mM NaCl; 3) 0.25 M LiCl, 1% Igepal, 1 mM EDTA, 10 mM TrisCl, pH8, 1% Deoxycholic acid. DNA protein complexes were eluted by two consecutive 15 min incubations at 65^0^C using elution buffer (0.1M NaHCO3, 1% SDS, 5mM DTT). The eluates were combined and crosslinking was reversed at 65^0^C for 5 hours. The samples were deproteinized and cleaned up with MinElute PCR purification kit (QIAGEN).

The sequencing library was prepared as previously described^9^ with minor modifications. Briefly: the end repair step was done in 1X T4 DNA ligase buffer with 10mM ATP in the presence of 0.25mM dNTPs, 0.6U T4 DNA polymerase, 0.5U Klenow Enzyme, 2 Units T4 Polynucleotide kinase for 30 min at 20^0^C. DNA was purified with MinElute kit. The second step was done in the presence of 1mM dATP and 1U of Klenow Exo minus. Reaction was incubated at 37^0^C for 30 min and DNA was purified with MinElute kit. To enrich for single stranded DNA the sample was denatured for 2 min at 95^0^C, then cooled to room temperature. The sequencing adaptor mix (Illumina) was diluted 1:200 and added for the ligation step. DNA was purified by MinElute kit and amplified using Phusion Polymerase (0.5 μl per reaction), in the presence of 1 μl of each Illumina PE primers. The following parameters were used: initial denaturation at 98°C for 30 sec; 21 cycles: 98°C 10 s, 65°C 30s, 72°C 30s; final extension for 5 min at 72°C. Size selection was done in 2% agarose gel, 180-250bp slice was excised and purified using MinElute Gel Extraction kit.

For H3K4me3 ChIP-Seq in oocytes, we isolated SCP3 positive meiotic oocytes from 14 15.5dpc females using BiTS-CHiP^54^. Briefly, this method uses FACS to isolate nuclei on the basis of the presence of an intra-nuclear marker (in this case, anti-SCP3 (Santa Cruz: sc-74569)). Kapa Hyper Prep kit (catalog #KR0961) was used to prepare the sequencing library due to the limited starting material relative to experiments in whole testis.

Testis sample preparation was performed as described previously^9^. The following antibodies were used: anti-DMC1: Santa Cruz (C-20, sc-8973), anti-DMC1: (custom), anti-H3K4me3: Millipore (#07-473). All sequencing was performed on an Illumina HiSeq 2500 at the NIDDK genomics core.

### Alignment of sequencing reads

For SSDS, reads were aligned to the genome and ssDNA derived reads were identified using the single stranded DNA sequencing (SSDS) processing pipeline^8^. Briefly, the first read of each mate pair is mapped to the genome with bwa (v0.7.12)^55^. The second read is then mapped to the genome using a modified bwa algorithm that finds the longest mapping suffix for each read. ssDNA is determined from the structure of inverted terminal repeats on the first and second end reads. The SSDS alignment pipeline is available at github^56^.

All other sequencing data were aligned to the reference genome using bwa aln (0.7.12)^55^.

### Evaluation of SSDS sensitivity for DSB detection in ovary

To estimate the lower hotspot detection limit using SSDS, we generated a DSB map using 2 x 10^5^ testis cells from Hop2^−/−^ (Psmc3ip^−/−^) mice^57^. HOP2 is required for DSB repair, and ~37% of cells in testis of Hop2^−/−^ mice harbor unrepaired DSBs (data not shown). The fraction of reads in peaks (FRiP) for this sample was 21% (Sample N1; Fig. S1A); this is slightly lower than the FRiP for DSB maps from whole testis (23-46%; Samples T1-9; Fig. S1A), but far above the 2% FRiP expected by chance. The estimated library size (see methods) of N1 was about 10-times smaller than for the smallest whole testis sample (Fig. S1D), likely because of the limited starting amount of DNA. Small library size complicates hotspot detection, therefore we first attempted to map DSBs in females using approximately 10 times more target cells (a pool of 90 ovaries; approximately 9 x 10^5^ target cells). This sample (O2) had a library size close to that of wt whole testis samples (Fig. S1D) but a lower FRiP (7%; Fig. S1A) suggested that hotspot DNA recovery from oocytes was less efficient than from testis. Subsequently, we pooled 230 fetal ovaries to generate a second ovary-derived DSB map (O1). This map was of similar quality to DSB maps derived from testis (FRiP = 33%; library size = 3.8 x 10^7^ fragments; Sample O1; Figure 1, Fig. S1A,D).

### DSB hotspot identification

Uniquely mapping fragments unambiguously derived from ssDNA (ssDNA type 1) and having both reads with a mapping quality score ≥ 30 were used for identifying hotspot locations (peak calling). NCIS^58^ was used to estimate the background fraction for each library. Peak calling was performed using MACS (v.2.1.0.20150420)^59^ with the following parameters : --ratio [output from NCIS]-g mm --bw 1000 --keep-dup all --slocal 5000. We use a mixture-model-based approach that accounts for GC-biases to calculate a corrected p-value for each hotspot (model = negative binomial; num. iterations for refinement = 100)^60^. P-values were adjusted for multiple testing using the Benjamini-Hochberg method. Hotspots with a GC-corrected P-value > 0.05 and DSB hotspots within regions previously blacklisted^10^ were discarded.

### H3K4me3 peak calling

Uniquely mapping reads with a mapping quality score ≥ 30 were used for peak calling. NCIS was used to estimate the background fraction relative to an input DNA library. Peak calling was performed using MACS (v.2.1.0.20150420)^59^ with the following parameters : --ratio [output from NCIS]-g mm --bw 1000 --keep-dup all --slocal 5000. Peak strength was subsequently calculated by subtracting the NCIS^58^ normalized input read count from the ChIP-Seq read count.

### Hotspot overlaps and merging hotspot sets

Unless otherwise stated, when assessing if hotspots occur at the same location, we restrict the overlap to the ± 200 bp region of DSB hotspots. Previously, we have shown that using a ± 200 bp regions is sufficient for detecting true overlaps and limits the number of spurious overlaps^9^. We merged DSB hotspots from the best testis and ovary samples (T1 and O1 respectively). The center of overlapping hotspots was defined as the mean center point of the T1 and O1 hotspot and the flanks were defined as the maximum distance from this center to the original T1 and O1 hotspot edges. Non-overlapping hotspots from each sample were retained as originally defined.

### Strength metrics at hotspots

Hotspot strength measured by SSDS was calculated as described previously^10^. Briefly, the center-point of the Watson- and Crick-strand ssDNA fragment distributions was used to define the hotspot center. ssDNA fragments on the Watson (top) strand to the left and Crick (bottom) strand to the right of this center were considered signal. Background was extrapolated from the count of ssDNA fragments of opposite polarity around the center (excluding the very center of the hotspot). Hotspot strength is then calculated as the signal - background fragment count. This strength is used as a proxy for DSB frequency^9,10^. Scripts for peak calling and strength quantitation are available at github^61^. Hotspot strength measured from Spo11-oligo mapping was calculated as the sum of the strengths of SPO11 oligo peaks overlapping each hotspot. SPO11-oligo peaks were downloaded from the processed data associated with the GEO record (GSM2247727)^21^. H3K4me3 strength at hotspots was calculated as the sum of the strength of overlapping H3K4me3 peaks.

### Sample FRiP reduction

FRiP is calculated as the number of in-hotspots ssDNA fragments divided by the total number of ssDNA fragments. To reduce FRiP, the required number of randomly selected in-hotspot fragments are discarded.

### Default hotspots

Hotspots that overlap a hotspot in mice that lack functional *Prdm9* were categorized as putative “default” hotspots.

### Sex bias determination

We used MAnorm^23^ to infer differential usage of DSB hotspots between the T1 and O1 samples. All hotspots from the T1, O1 merge were used as “common” peaks. Hotspots on chromosomes X, Y and M were excluded from this analysis. MAnorm p-values were adjusted for multiple testing using the Benjamini-Hochberg method and hotspots with a corrected p-value < 0.01 were considered differential.

### Cluster analysis

We identified groups of adjacent hotspots that shared the same sex-bias (female-biased, male-biased, unbiased). Only uninterrupted runs of hotspots with the same bias were considered. To estimate the expected numbers of clusters, hotspot bias designations were shuffled and the aforementioned process was repeated. 10,000 iterations of this randomization process were performed. P-values are calculated from the empirical distribution of expected values for clusters of each size.

### Generation of randomized maps of DSB hotspots

DSB hotspot locations were randomized as described in^9^. Briefly, hotspots were uniformly distributed per chromosome, but prohibited from being placed at unmappable regions. Specifically, a mapability score for 40bp sequencing reads at each mm10 base was calculated using the GEM library (20100419-003425)^62^. Hotspot excluded regions were defined as annotated assembly gaps 9from UCSC)^63^, in addition to 1 kb genomic intervals with <50% uniquely mappable bases plus 1 kb either side. Hotspot width and strength were preserved at the randomized location for each hotspot.

### Genetic crossovers

The locations of mouse crossovers in were obtained from the Collaborative Cross^33^. To assess if the elevated male crossover rate in the q-arm subtelomeric region is *Prdm9* dependent, we inferred genetic crossovers that were likely to have been formed by the B6, CAST or PWD alleles of PRDM9. Crossovers likely defined by the B6, CAST and PWD/PWK alleles of PRDM9 were determined by first identifying crossovers that occurred in hybrids involving any of these pure strains (MGP,MGM,PGP,PGM). Crossovers that coincided with a single DSB hotspot from only one of the parental PRDM9 alleles were then designated as having originated from that allele. All hotspots between B6, 129 and WSB mice that coincided with a B6, C3H or 13R-defined DSB hotspot were designated as having originated from *Mus musculus domesticus* (Dom). COs that did not overlap any DSB hotspot from either parental genotype and COs from crosses for which DSB hotspot maps were not available were designated as “Ambiguous”. Crossovers derived from all PRDM9 alleles showed similar sub-telomeric enrichment for males relative to females, consistent with previous reports^64^. Thus, this enrichment appears a Prdm9-independent phenomenon.

### DNA methylation

For BS_SC_, BS_SG_, BS_DAA_, BS_DWT_ (Table S1), bismark^65^ was used to align whole genome bisulfite sequencing (WGBS-Seq) data to the reference mm10 genome. For BSrahu, BS_PGC_, BS_LIV_, BS_MG_ (Table S1), pre-processed nucleotide-resolution methylation data were available, therefore UCSC liftover^63^ was used to convert these data from mouse mm9 to mouse mm10 genomic coordinates where necessary. hMeDIP-Seq reads^43^ were mapped to the genome using bwa mem (0.7.12)^55^ and default parameters.

To examine methylation patterns, we first inferred a high confidence set of PrBSs at DSB hotspots. We used FIMO^66^ to identify matches to the B6 PrBS9 PWM within hotspots. The number of matches to the position weight matrix (PWM) of the PrBS depends on the alignment score threshold used, therefore we identified the PWM p-value alignment threshold that yielded the maximal number of DSB hotspots with a single match to the PRDM9 PWM within the central ± 250 bp. We tested the following threshold values: P ≤ 5x10^−3^, 1x10^−4^, 2x10^−4^, 4x10^−4^, 6x10^−4^, 8x10^−4^, 1x10^−5^, 2x10^−5^, 4x10^−5^ 4x10^−5^, 6x10^−5^, 8x10^−5^, 8x10^−5^,1x10^−6^, 1x10^−7^, 1x10^−8^. 12,097 / 19,053 hotspots (63%) contained a single PrBS at the optimal threshold (P ≤ 4x10^−4^).

### High copy repeats at DSB hotspots

High copy repeats determined by repeatmasker^67^ for the human hg38 genome were split into 67 groups by family. Repeat families that overlapped < 0.2% of any hotspot set were excluded. We used UCSC liftOver^63^ to convert DSB hotspots from the hg19 to hg38 genome coordinates. To assess if hotspots biased to each sex associated with different DNA high copy repeat families, we counted the frequency of repeats overlapping the central ± 200 bp of female-biased, unbiased and male-biased hotspots. To assess differences, we performed a two-sided binomial test for all pairwise comparisons (male/female, male/unbiased, female/unbiased). P-values were Bonferroni corrected to account for multiple testing and a corrected P-value of P < 0.01 was used to assess differences. Repeats that showed significantly different enrichment in one set of hotspots compared to both others were investigated. From this analysis, two families of LTR retrotransposons (LTR & LTR/ERVK) are depleted at male-biased hotspots, whereas LINE/L1, SINE/Alu and SINE/1D elements are depleted at female-biased hotspots relative to the other two sets. Notably however, no repeat families are elevated exclusively at either set of sex-biased hotspots.

## Supplementary Data

**Supplementary Data Figure 1:**
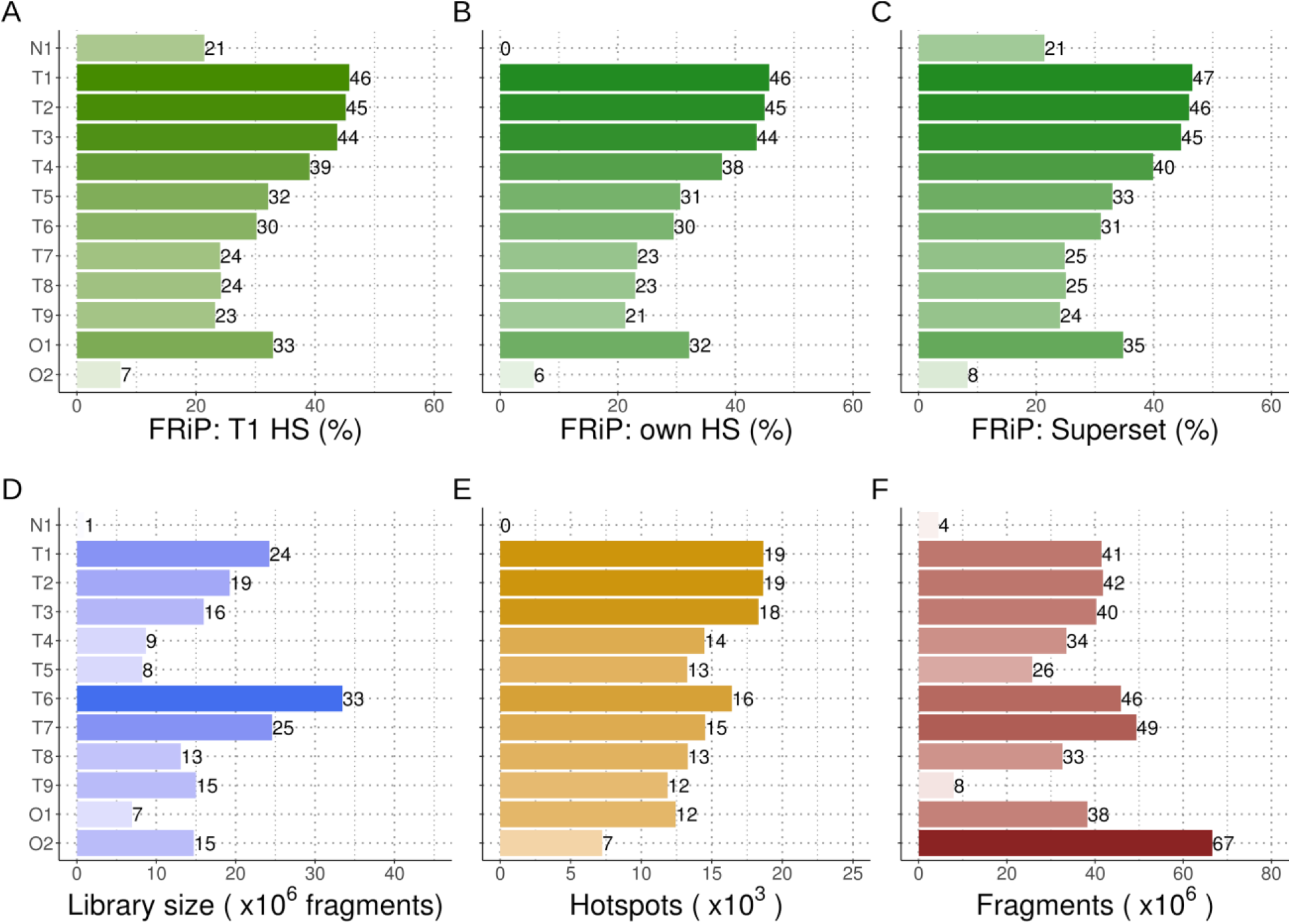
Sample details and quality metrics for DSB maps. The fraction of reads in peaks is calculated for all samples at (A) hotspots identified in the T1 sample. For sample identifiers, see Table S1. (B) Hotspots identified in each respective sample or (C) hotspots in the combined T1/O1 superset. Peak calling was not performed for N1 (see methods). (D) The estimated library size (x) was inferred using bisection root finding for f(x) = (1-N_NR_/x)-exp(N_TOT_/x), 10^4^ <= x <= 10; N_NR_ = # unique fragments; N_TOT_ = # total fragments. (E) The number of hotspots identified in each sample. (F) The number of ssDNA fragments sequenced for each sample.

**Supplementary Data Figure 2:**
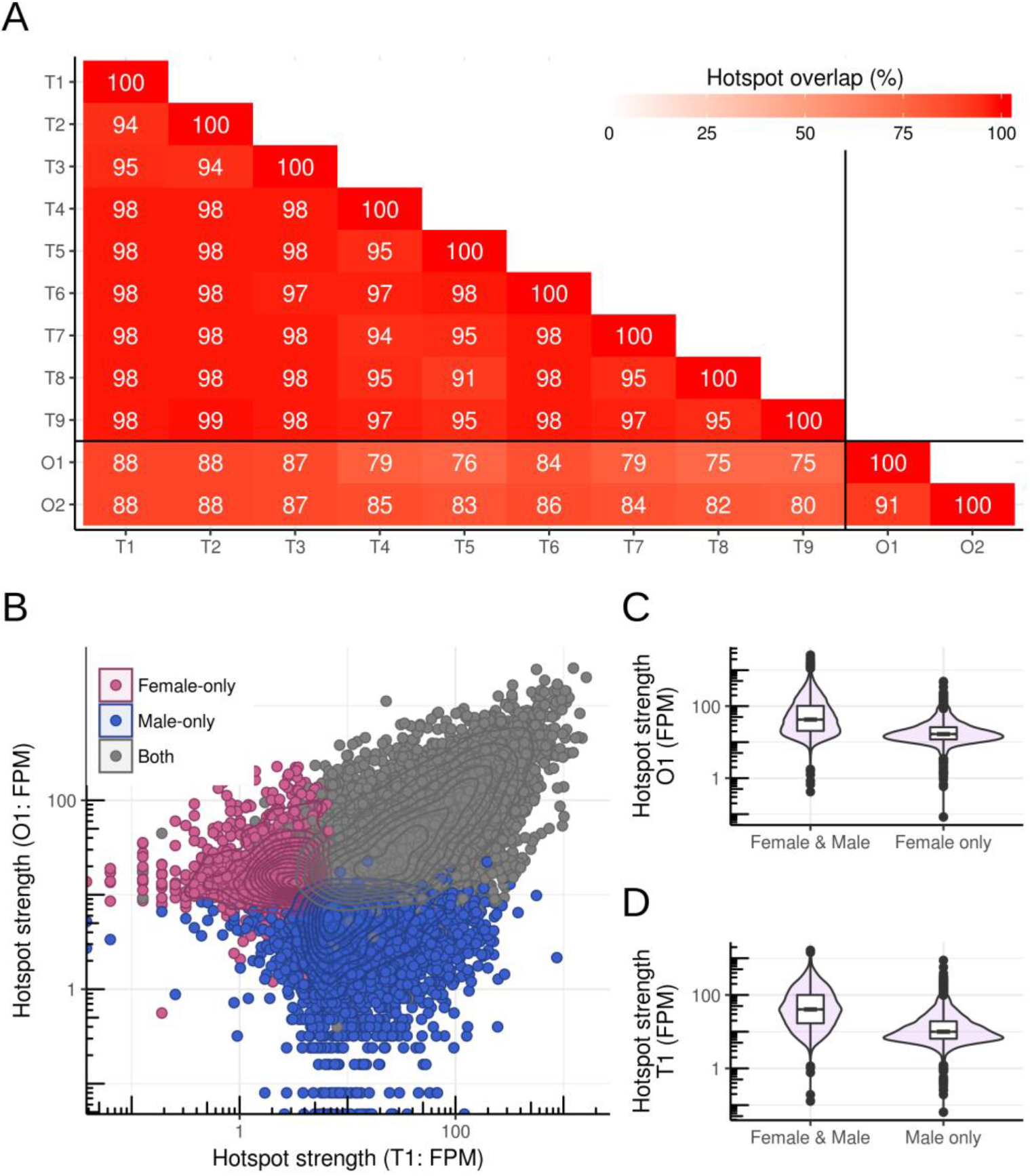
Most DSB hotspots are used in both male and female meiosis. (A) The overlap between hotspots in each sample was calculated using only the central ± 200 bp of hotspots. The maximum reciprocal overlap between samples is shown. As shown in Extended Figure 1, the overlap with O2 is low, partly because of spurious peak calls that were not removed for this analysis. (B) Hotspots exclusively found in either sex are weak. Hotspots were split into those found in O1 and T1 (Both; grey), O1, not T1 (Female-only; pink) and T1, not O1 (Male-only; blue). (C) Female-only hotspots are weak in females, relative to shared hotspots. (D) Male-only hotspots are weak in males, relative to shared hotspots.

**Supplementary Data Figure 3:**
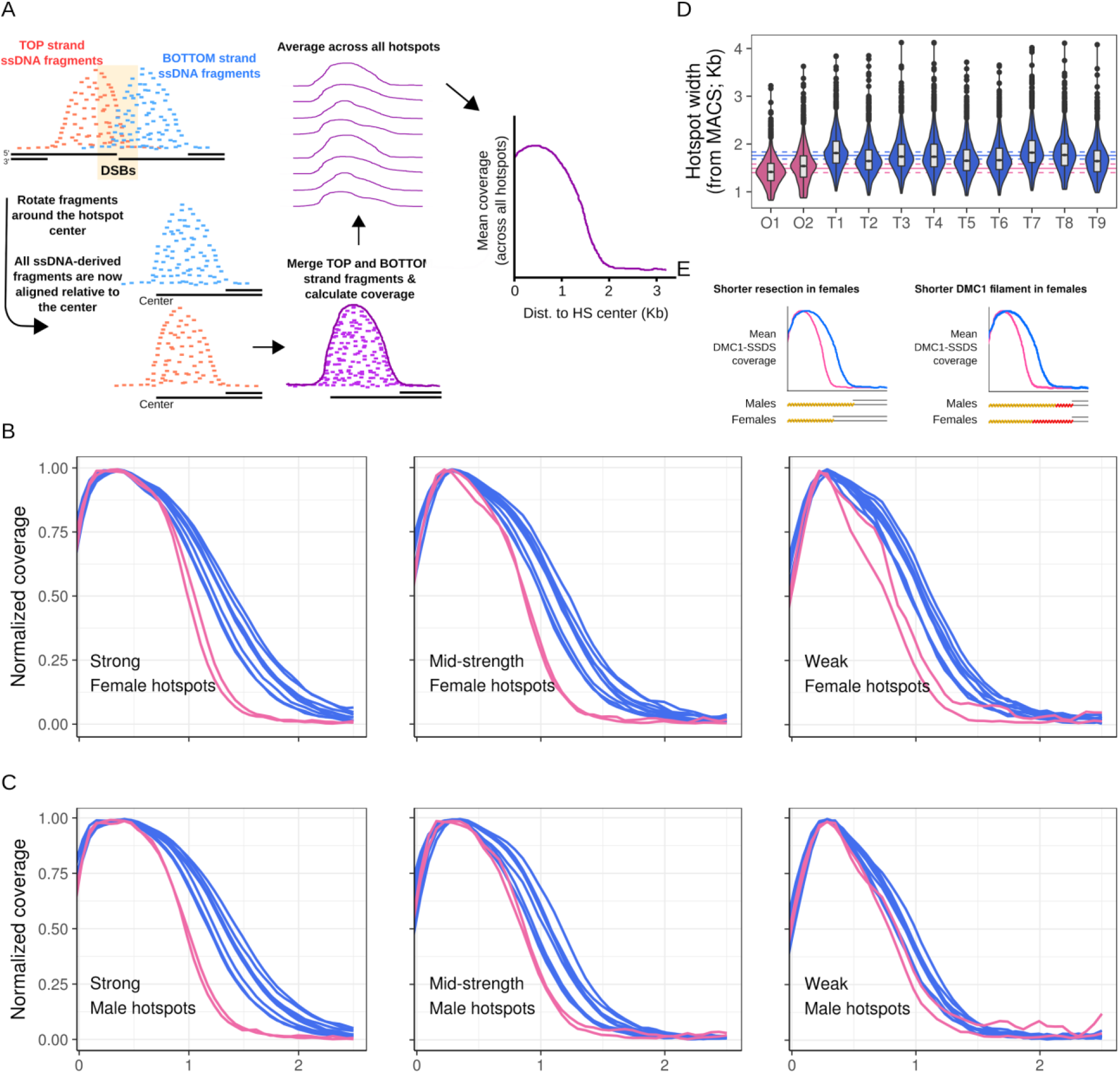
SSDS signal at hotspots is narrower in ovaries than in spermatocytes. (A) SSDS coverage is a measure of DMC1-bound ssDNA either side of each meiotic DSB. In a population of meiocytes, DSBs will occur in a several hundred nucleotide window around the hotspot center (orange rectangle). To assess coverage, we first convert the position of each SSDS fragment into the distance along ssDNA from the hotspot center. Merging the top and bottom strand fragments in this way increases coverage two-fold and minimizes the influence of asymmetric gaps and fluctuations in coverage. Coverage at each hotspot was normalized by the maximum value at the hotspot to prevent strong hotspots from dominating the average profile. The average normalized coverage across all hotspots was then calculated. DSB hotspots identified in (B) females (ovary sample O1) and (C) males (testis sample T1) were each split into three bins by strength. Coverage was calculated for all nine male and two female samples for each set. The SSDS signal is narrower for all female samples compared to male samples. The difference is particularly pronounced at stronger hotspots, where coverage estimates are most accurate. At the widest point, the mean male and female profiles diverge by ~0.4 Kb. (D) We also examined the MACS-determined hotspot boundaries to further negate the possibility that the average profiles in B-C are not a reflection of the population. By this metric, the mean hotspot width estimated from male samples (1,759 ± 73 bp; mean (solid blue line) ± SEM (dashed blue lines); n = 9) is significantly wider the mean width of hotspots in female samples (1,490 ± 89 bp; mean (solid pink line) ± SEM (dashed pink lines); n = 2) (t-test; P = 0.0007). Sequencing quality and sample FRiP can affect width estimates, therefore, we processed each sample as follows; we reduced the FRiP of each sample to that of the lowest quality sample (O2; see methods), considering only uniquely mapping and high quality (Q>30) ssDNA type 1 fragments. We then reduced all samples to have the same number of fragments as the smallest. On these datasets, we performed peak calling and retained only DSB hotspots that were called in all samples (N = 1,975). (E) Potential mechanistic explanations for the difference in SSDS signal between males and females. These differences may manifest in all meiocytes or in sub-populations. Importantly, we see no evidence of shape differences at hotspots in sub-populations of spermatocytes (data not shown).

**Supplementary Data Figure 4:**
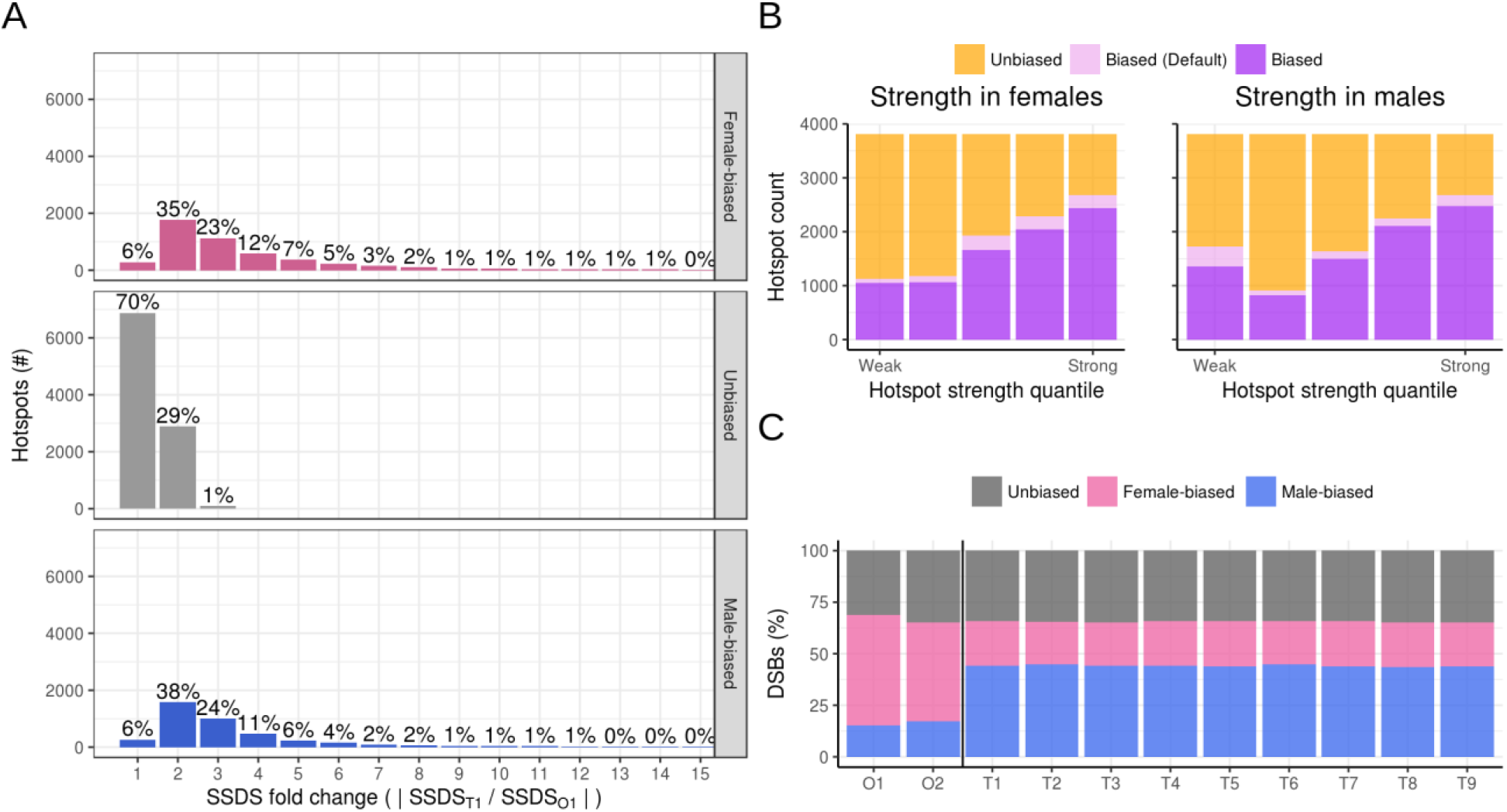
Most meiotic DSBs occur at sex-biased hotspots. (A) Quantification of SSDS fold change at sex biased and unbiased hotspots. The percentages show the percentage of hotspots in each category with a given absolute fold-change. (B) The hotspots in the testis/ovary super set were split into quintiles by strength in either females (left panel) or males (right panel). In both sexes, over 60% of the strongest hotspot subset exhibit sex-biased DSB formation. This is a proxy for the true amount of sex-biased DSB formation. In progressively weaker hotspot sets, fewer biased hotspots are detected. One outlier is the set of weak male hotspots. This set contains many female-biased default hotspots that form independently of PRDM9. (C) We quantified the total in-hotspot SSDS signal at female-biased, unbiased and male-biased hotspots in the two ovary-derived samples and in the nine testis samples. In all cases, over half of the in-hotspot sequencing tags (referred to as total DSBs) occur at sex-biased hotspots. Hotspots biased towards usage in females are enriched in ovary samples, while those biased towards male usage are enriched in testis-derived samples.

**Supplementary Data Figure 5:**
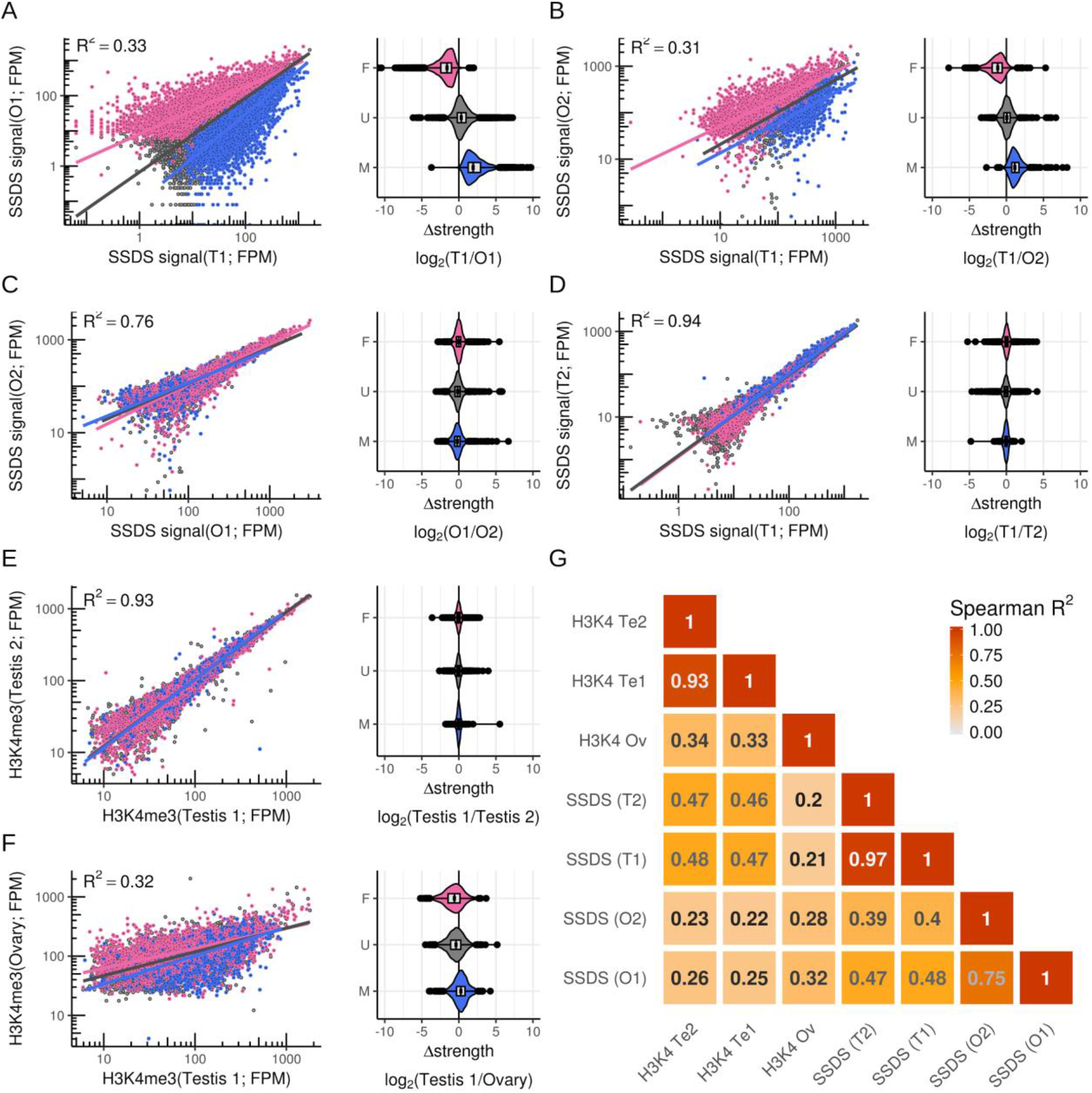
Sex biased hotspots are consistent across replicates and are defined before DSB formation. Hotspot strength was calculated at all autosomal hotspots from the merged O1/T1 DSB maps. The strength of hotspots was re-calculated in two testis (T1, T2) and two ovary (O1, O2) maps. Female-biased (pink), unbiased (grey) and male-biased hotspots (blue) were determined by comparing the T1 and O1 maps. These hotspots are colored the same in all panels. (A) Sex-biased hotspots are distributed as expected when comparing the O1 and T1 DSB maps. These data are also plotted in Fig. 2D,E. (B) Sex-biased hotspots exhibit the same sex-biases in the O2 sample. (C,D) Sex-biased hotspots exhibit no biased usage between samples derived from mice of the same sex. (E-G) Sex biases that precede DSB formation were studied by performing H3K4me3 ChIP-Seq in fluorescence assisted cell sorting (FACS)-purified fetal oocytes at 15.5 dpc (see methods). The H3K4me3 signal at hotspots was quantified and compared to existing maps of H3K4me3 in juvenile mouse testis^<u>68</u>^. (A) The H3K4me3 signal at hotspots is tightly correlated in replicate samples from mouse testis. (B) Akin to what we observe when examining the SSDS signal at DSB hotspots, there is extensive variation in the H3K4me3 signal at hotspots between male and female meiosis. This implies that sex biases are established before DSB formation. Sex biases determined using SSDS remain broadly conserved when we compare H3K4me3 in females to males. (C) H3K4me3 at hotspots is better correlated with SSDS from the respective sex.

**Supplementary Data Figure 6:**
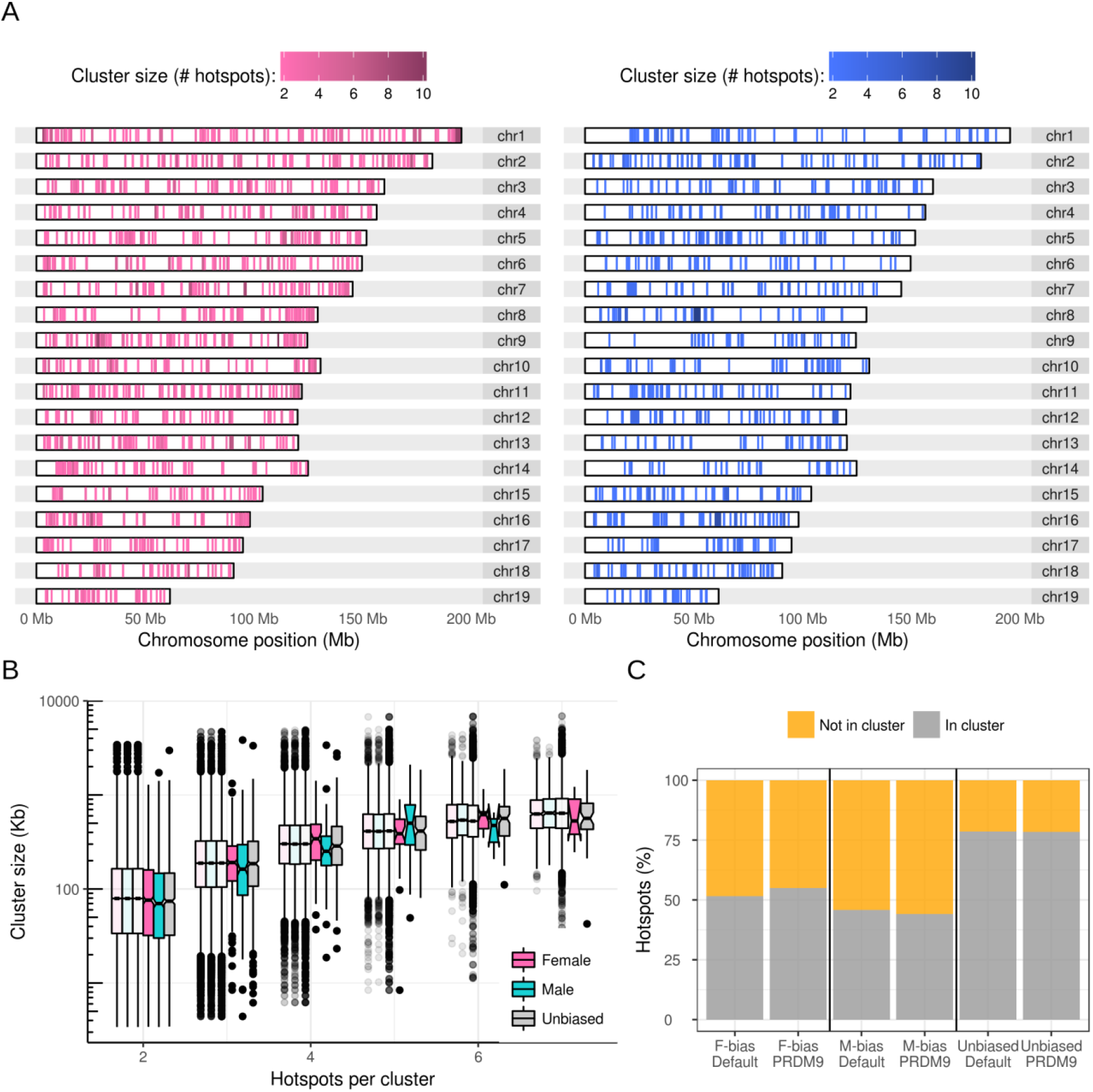
Clustering of sex-biased hotspots. (A) Genomic patterning of sex-biased DSB hotspots. Female-biased (left; pink) and ale-biased (right; blue) hotspot clusters on all autosomes. Biased hotspots do not exhibit particular spatial patterning, aside from a slight enrichment of female-biased hotspots at the q-Arm telomere. (B) The physical size of hotspot clusters scales with the number of hotspots per cluster. It therefore seems unlikely that clustering results from a physical size constraint imposed by sex-specific chromatin structure. Importantly however, the presence of such a size constraint may be masked by the presence of a large number of clusters that occur by chance. Semi-transparent boxplots show the expected size distribution for randomly distributed clusters (n = 1,000 bootstraps). Clusters of three male-biased hotspots are marginally smaller than expected. There are no significant differences for clusters of other sizes. (C) Similar proportions of PRDM9-defined and default hotspots occur in clusters. Hotspots in clusters of ≥ 2 consecutive hotspots of the same type were counted.

**Supplementary Data Figure 7:**
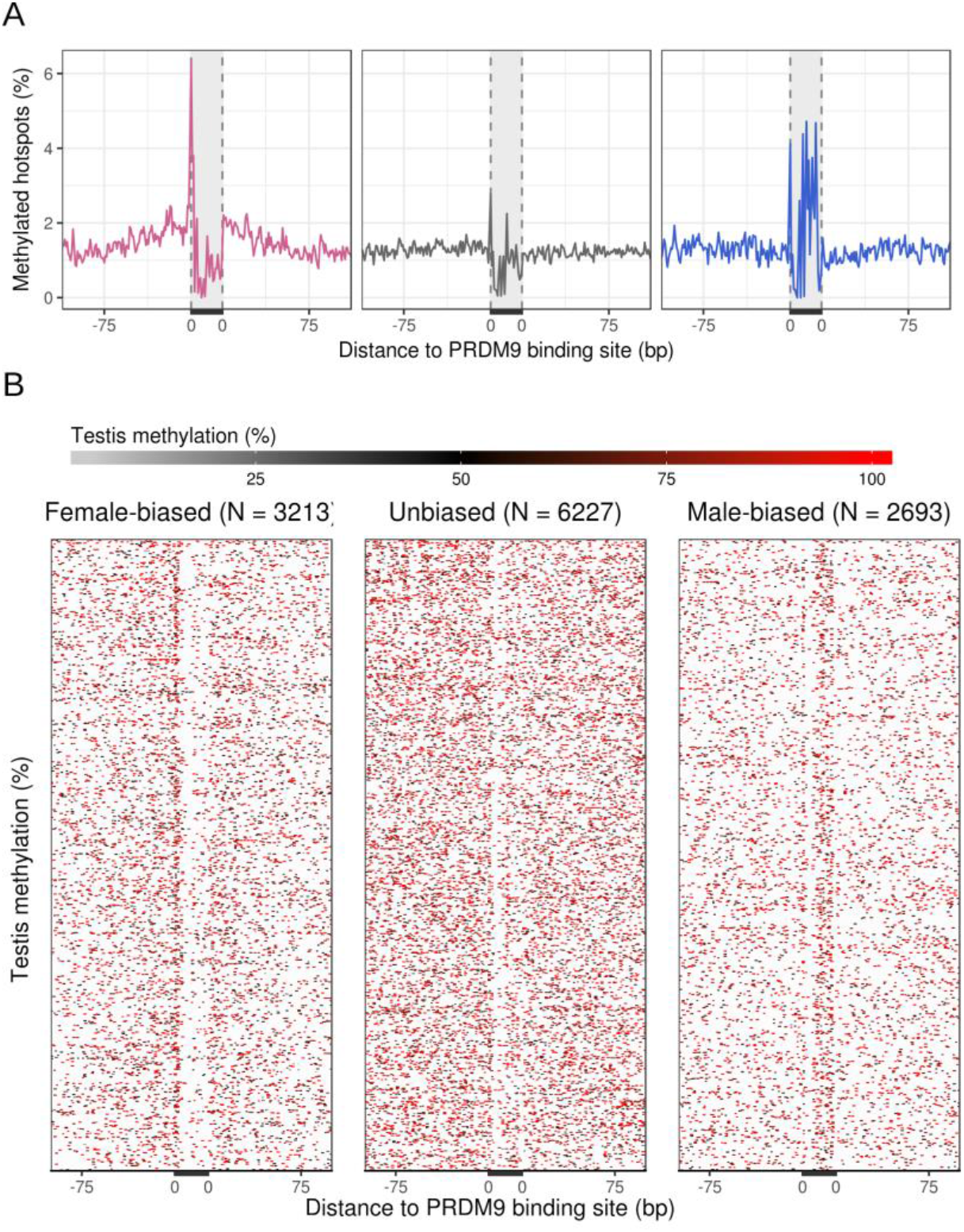
Differing patterns of DNA methylation at sex-biased hotspots. (A) Mean DNA methylation^40^ at the putative PrBS (grey bar) of female-biased (pink), unbiased (grey) and male-biased (blue) hotspots. Note that this panel is also shown in Figure 4A. (B) Heatmap rows depict methylation at individual hotspots. Note that the density of methylation appears higher at unbiased hotspots because rows are more densely spaced.

**Supplementary Data Figure 8:**
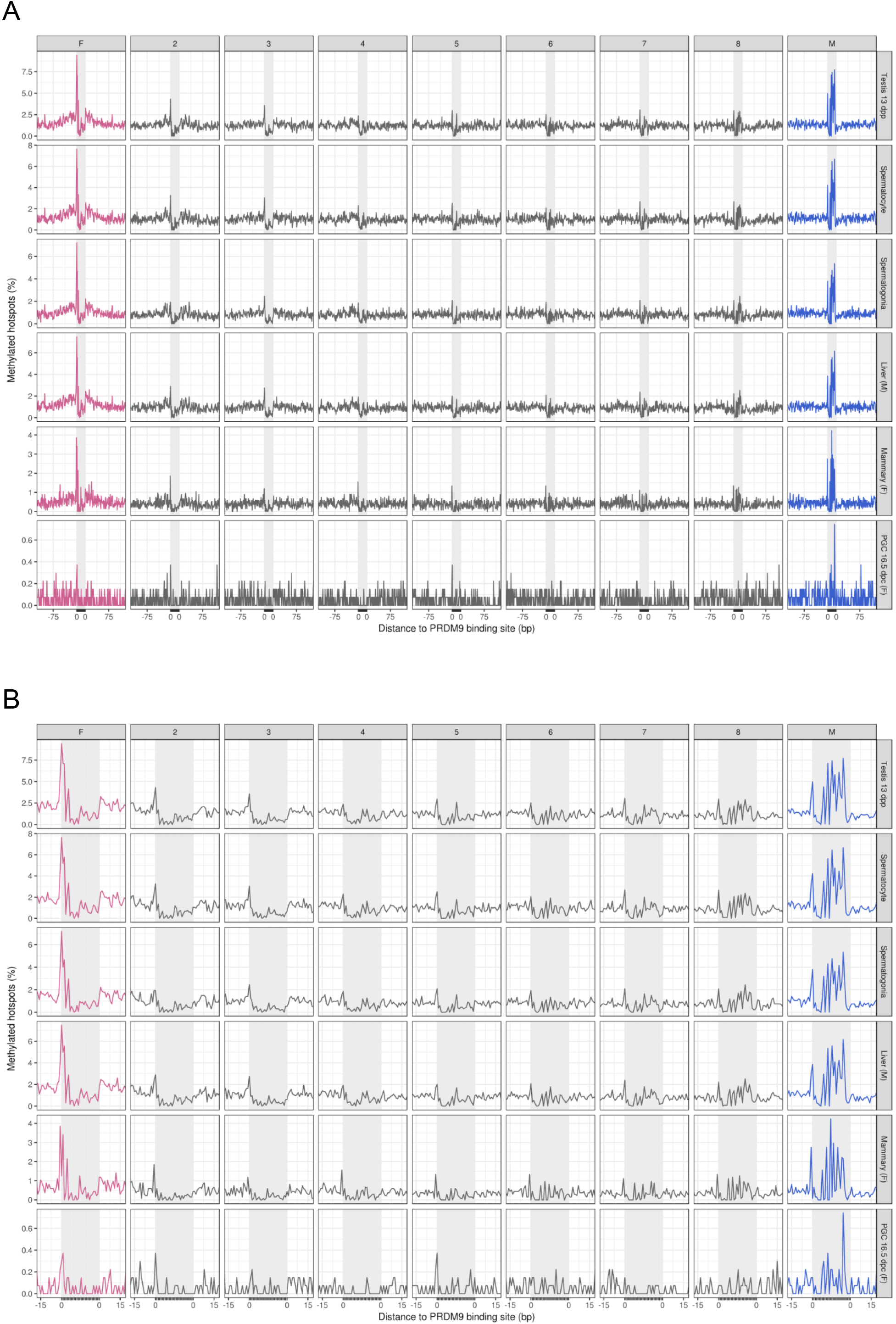
DNA methylation at PrBSs is present across tissues and absent in the female germ line. The pattern of DNA methylation is very similar across cell types and between the sexes. Hotspots are split by the magnitude of sex bias (SSDS_O1_/SSDS_T1_) into nine sets. Sets are ranked from most female-biased (pink; left) to most male-biased (blue; right) by fold-change. Methylation signal is binarized such that methylation > 0% is considered methylated. Thus, the proportion of all hotspots with methylated cytosine at each position is shown. Variations in the magnitude of the signal may be expected for technical reasons. Plots are anchored by the C57B6 PrBS (grey area). (A) Plot of ± 100 bp to show methylation flanking the PrBS for female-biased hotspots. (B) Plot of ± 15 bp to show methylation at the PrBS for male-biased hotspots. Methylation data are from whole genome bisulfite sequencing experiments in tissue derived from whole testis in 13dpp mice^40^, elutriated spermatocytes^43^, spermatids^43^, male liver^42^, female parous basal differentiated mammary gland cells^41^, and sorted primordial germ cells at 16.5 dpc^36^.

**Supplementary Data Figure 9:**
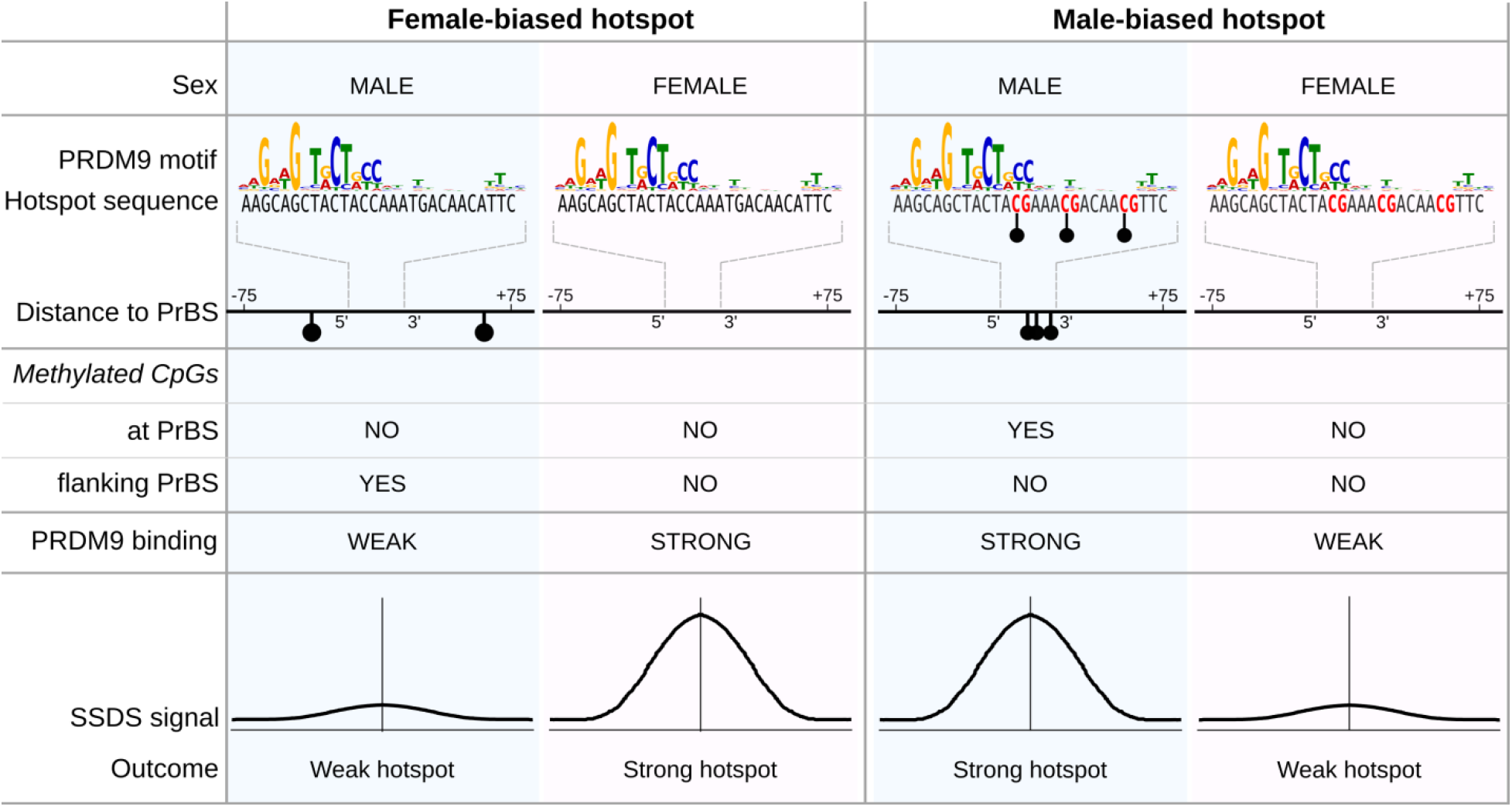
Dual role of DNA methylation at hotspots in defining sex biases. DNA methylation has a dual role in modulating sex-biased DSB formation. (Left panels) At female-biased hotspots, DNA methylation in the region flanking the PrBS can suppress PRDM9 binding. Thus, in males, the use of these PrBS is reduced, resulting in a female-biased hotpsot. Methylated CpG dinucleotides (in males) are schematically shown as filled black circles. (Right panels) At male-biased hotspots, DNA methylation at CpGs appears to favor PRDM9 binding and DSB formation. This results in a relatively strong DSB hotspot in males, but a relatively weak hotspot in females, where DNA methylation at these sites is absent.

**Supplementary Data Figure 10:**
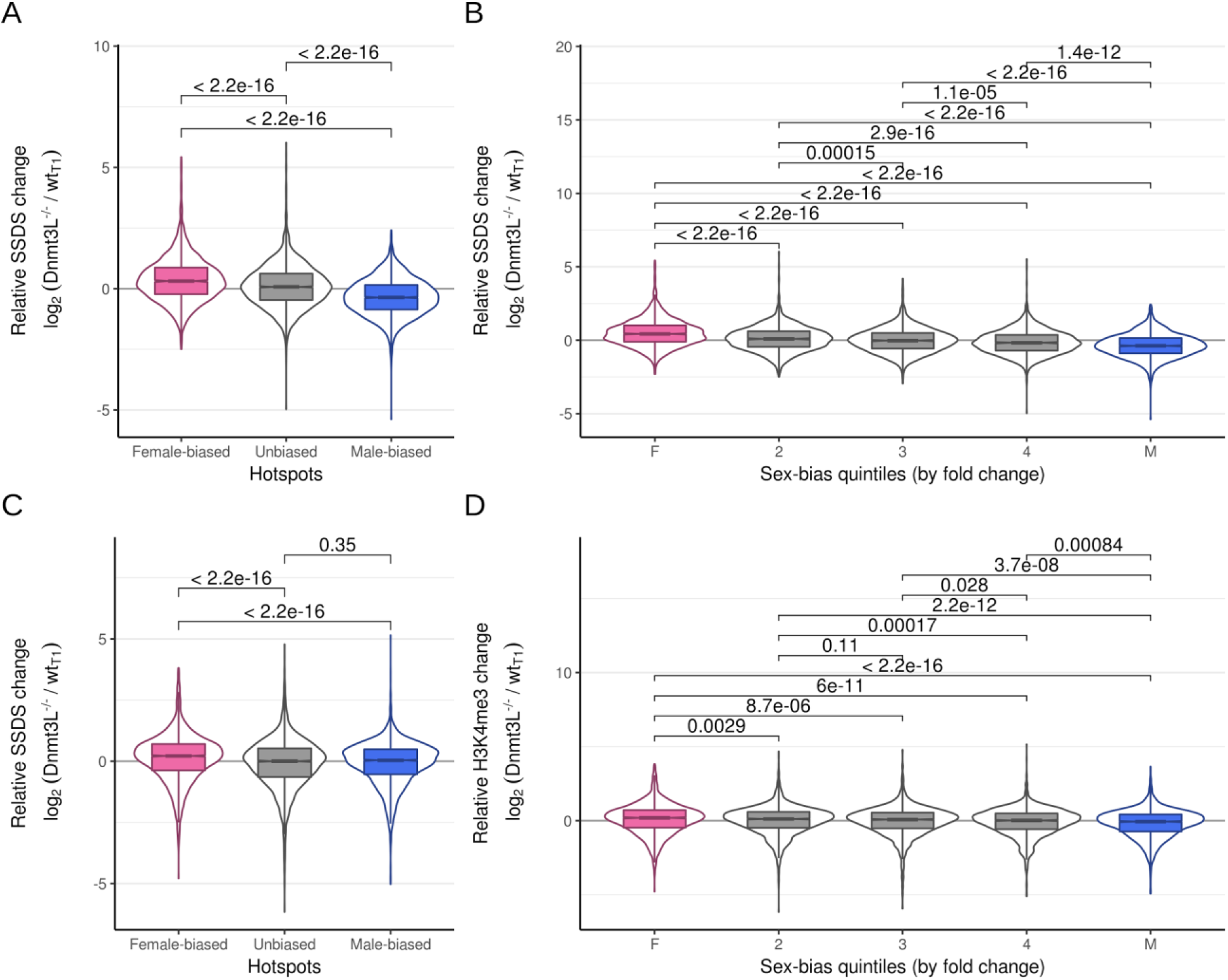
Hotspot strength variation in Dnmt3L^−/−^ mice. The usage of sex-biased hotspots is altered in mice where DNA methylation is reduced (Dnmt3L^−/−^). Hotspots are split by assigned sex bias (A,C) or into quintiles by the fold change between O1 and T1 SSDS samples (B,D). The log2 fold change between the tags per million normalized signal at hotspots in Dnmt3^−/−^ and wt (T1) male mice is shown. Hotspots overlapping gene promoters or default hotspots are excluded as the non-PRDM9 derived H3K4me3 signal would confound this analysis. Furthermore, only hotspots detected in all samples being compared were analysed to remove spurious potential background correlation. P-values for all comparisons are shown above the data (wilcoxon test). (A) The SSDS signal at female-biased hotspots is significantly increased in Dnmt3L^−/−^ mice compared to male-biased hotspots or unbiased hotspots. The strength of male-biased hotspots is relatively decreased. (B) This is also seen when we simply split hotspots by fold change. (C) H3K4me3 at female-biased hotspots is significantly increased in Dnmt3L^−/−^ mice compared to male-biased hotspots. H3K4me3 signal at each hotspot was calculated as the sum of overlapping H3K4me3 peak strengths. This is a proxy for DSB hotspot strength, because PRDM9 trimethylates histone H4 lysine 3 before DSB formation. (D) This is more apparent when we split hotspots into quintiles by sex-bias, likely because H3K4me3 at hotspots is a weak signal.

---

**Supplementary Table S1:** SSDS sample details. Testis samples were named T1-9. Ovary samples were named O1,O2. Sample N1 was generated from *Hop2^−/−^* mice using 2 x 10^5^ cells. This sample was run in single-end mode, therefore it was not processed through the ssDNA pipeline. Samples that have been previously published are referenced by their GEO accession number.

| Sample | Source | Antibody | Genotype | Accession # | Start material |
| --- | --- | --- | --- | --- | --- |
| N1 | Testis (adult) | SC 8973 (c20) | B6 *Hop2*<sup>-/-</sup> | n/a | 2 x 10<sup>5</sup> cells |
| T1 | Testis (adult) | SC 8973 (c20) | B6 *wt* | GSM869781 | Whole testis |
| T2 | Testis (adult) | SC 8973 (c20) | B6 *wt* | n/a | Whole testis |
| T3 | Testis (adult) | SC 8973 (c20) | B6 *wt* | n/a | Whole testis |
| T4 | Testis (adult) | SC 8973 (c20) | B6 *wt* | n/a | Whole testis |
| T5 | Testis (adult) | SC 8973 (c20) | B6 *wt* | n/a | Whole testis |
| T6 | Testis (adult) | SC 8973 (c20) | B6 *wt* | n/a | Whole testis |
| T7 | Testis (adult) | αDMC1-GP1 | B6 *wt* | n/a | Whole testis |
| T8 | Testis (adult) | SC 8973 (c20) | B6 *wt* | n/a | Whole testis |
| T9 | Testis (adult) | SC 8973 (c20) | B6 *wt* | GSM869782 | Whole testis |
| O1 | Ovary (15.5 dpc) | αDMC1-GP1 | B6 *wt* | n/a | 230 ovaries |
| O2 | Ovary (15.5 dpc) | αDMC1-GP1 | B6 *wt* | n/a | 90 ovaries |
| TP1 | Testis (adult) | SC 8973 (c20) | *Prdm9*<sup>-/-</sup> | GSM869796 | Whole testis |
| DN1 | Testis (30 dpp) | SC 8973 (c20) | *Dnmt3L*<sup>-/-</sup> | n/a | Whole testis |
| iO1 | Ovary (15.5 dpc) | n/a : input | B6 *wt* | n/a | 230 ovaries |

**Supplementary Table S2:** Other sequencing sample details

| ID | Sample | Source | Antibody | Genotype |
| --- | --- | --- | --- | --- |
| H3<sub>W30</sub> | H3K4me3 ChIP-Seq | Testis (30 dpp) | Millipore (#07-473) | B6 *wt* |
| H3<sub>D30</sub> | H3K4me3 ChIP-Seq | Testis (30 dpp) | Millipore (#07-473) | *Dnmt3L*<sup>-/-</sup> |

**Supplementary Table S3:** Public datasets used

| ID | Sample | Data accession |
| --- | --- | --- |
| T1 | C57Bl/6 SSDS from testis | GSM869781 |
| T9 | C57Bl/6 SSDS from testis | GSM869782 |
| PR | *Prdm9*<sup>-/-</sup> SSDS from testis | GSM869790 |
| H3<sub>H2</sub> | C57Bl/6 Hop2<sup>-/-</sup> H3K4me3 ChIP-Seq | GSM869791 |
| SPO | Spo11-associated oligo mapping from C57Bl/6 mice | GSM2247727 |
| BS<sub>RA</sub> | C57BL/6J & FVB/NJ whole genome bisulfite sequencing (WGBS) from 13 dpp male testis | GSM2698071 |
| BS<sub>SC</sub> | C57Bl/6 WGBS from sorted spermatocytes | GSM1202742, GSM1202743 |
| BS<sub>SG</sub> | C57Bl/6 WGBS from sorted spermatogonia | GSM1202738, GSM1202739 |
| BS<sub>MG</sub> | Balb/C WGBS from parous basal differentiated cells in female mammary gland | GSM1646791 |
| BS<sub>LIV</sub> | C57Bl/6 WGBS from male liver | SRR2173867 to SRR2173871 |
| BS<sub>PGC</sub> | C57Bl/6 WGBS from female E16.5 primordial germ cells | ERX167033 |
| hM<sub>SC</sub> | C57Bl/6 hMeDIP-Seq from spermatocytes | GSM1202717 |
| hM<sub>ST</sub> | C57Bl/6 hMeDIP-Seq from spermatids | GSM1202718 |
| hM<sub>SP</sub> | C57Bl/6 hMeDIP-Seq from sperm | GSM1202719 |
| hM<sub>LivM</sub> | hMeDIP-Seq from male liver | ERR1702166 |
| hM<sub>LivM</sub> | hMeDIP-Seq from female liver | ERR1702154 |
| hM<sub>LivM</sub> | hMeDIP-Seq from male cortex | ERR1702160 |
| hM<sub>LivM</sub> | hMeDIP-Seq from female cortex | ERR1702148 |

